# Thrombin cleavage of circulating osteopontin in the lung drives hippocampal neuroinflammation and acute delirium-like neurologic dysfunction after acute kidney injury

**DOI:** 10.64898/2026.11.17.767415

**Authors:** Poornima Dilhani Ekayanake Weerasinghe-Mudiyanselage, Rupal Ojha, Liang Ning, Marta Celorrio, Leah Tolby, Anusha Suresh, John Morser, Lawrence Leung, Machiko Honda, Shigeyuki Kon, Stuart H. Friess, Andreas Herrlich

## Abstract

Acute kidney injury (AKI) is a common clinical problem with high morbidity and mortality due to frequently occurring secondary organ complications, including acute neurologic dysfunction, such as delirium. Specific mechanisms responsible for this kidney-brain crosstalk are unknown but hypothesized to include uremic toxins and neuroinflammation secondary to systemic inflammation. Here we show that uremic mice with AKI show hippocampal monocytic neuroinflammation associated with acute delirium-like neurologic dysfunction only in the presence of circulating osteopontin fragments. Mechanistically, circulating osteopontin (OPN/Spp1) released by the injured kidney was cleaved by thrombin in the lung into OPN-N and OPN-C-terminal fragments. Both OPN fragments were required to drive upregulation of the monocyte chemoattractant chemokine-ligand-2 (CCL2) in hippocampal endothelial cells. AKI mice lacking OPN or unable to activate OPN by thrombin cleavage showed kidney injury and serum uremic toxin elevations like *wt* controls but were protected from hippocampal CCL2 upregulation, monocyte recruitment, and delirium-like symptoms. Systemic application of an OPN or CCL2 neutralizing antibody was protective. Our results suggest that a specific kidney-released circulating factor, osteopontin, drives acute neurologic dysfunction after AKI and challenge the widely held hypothesis that uremic toxins play a dominant role. They also identify thrombin cleavage of OPN as a potential therapeutic target and critical switch for the remote functions of circulating OPN.

## Introduction

Acute kidney injury (AKI) develops in 2-5% of hospitalized patients, and in 30-50% of critically ill patients^1–4^. Major AKI-induced morbidity and mortality occur largely due to secondary remote organ complications not treatable by dialysis^5^, including e.g. acute delirium^6^ and chronic neurocognitive dysfunction^7–9^, cardiovascular^10,11^ and pulmonary complications^12–15^. AKI acutely causes delirium in 60% of patients^16^, and chronically increases the risk of dementia and cognitive impairment^17^, significantly affecting brain health over the lifespan. How AKI negatively impacts the brain acutely or chronically is largely unknown.

Central hypotheses around the causes of AKI-induced acute delirium are largely based on correlative evidence obtained in chronic kidney disease (CKD) patients or CKD animal models and implicate uremic retention solutes (“uremic toxins”), circulating classical cytokines (e.g. tumor necrosis factor or interleukin-6), increased blood-brain-barrier (BBB) permeability, and neuroinflammation involving the hippocampus and cortex. However, mechanistic studies testing these hypotheses directly are largely missing^18–20^. Acute delirium after AKI has not been studied in pre-clinical models, but AKI studies in rodents indeed reported signs of acute neuroinflammation. Studies in mice at 24 hours after ischemia-reperfusion of the kidney showed cytokine and chemokine upregulation in cortex and hippocampus, microglial activation, and – in one study - decreased locomotor activity^6,21,22^, suggesting the existence of kidney-brain interorgan crosstalk after AKI. However, how AKI established neuroinflammation mechanistically, how this caused locomotor dysfunction, and whether neuroinflammation caused delirium was not studied.

We previously reported that circulating osteopontin (OPN/SPP1) released by the injured kidney after AKI causes remote lung inflammation and respiratory failure by acting on lung intravascular CCR2^+^ monocytes and their lung interstitial macrophage progeny^23,24^. Whether OPN is involved in other secondary organ complications, such as AKI-induced neuroinflammation and delirium is not known.

Here we show in mice that AKI induces acute hippocampal monocytic inflammation with delirium-like neurologic dysfunction that depends on thrombin cleavage of circulating OPN in the lung, but not on increases in blood brain barrier permeability, or solely on effects of uremic toxins. We define OPN fragment target cells in the hippocampus and show that OPN fragment-induced expression of the monocyte chemoattractant chemokine-ligand-2 (CCL2) in hippocampal endothelial cells drives AKI-induced hippocampal neuroinflammation. Results in our pre-clinical model of AKI-induced delirium suggest that neuroinflammation induced by a specific kidney-released circulating factor, osteopontin, is responsible for acute delirium-like neurologic dysfunction after AKI and at least question the widely held hypothesis that uremic toxins play a dominant role in this process.

## Results

### AKI induces acute neurological dysfunction with delirium-like symptoms in mice

Delirium is a common neuropsychiatric syndrome in particular in hospitalized patients. It is characterized by acute onset of deficits in attention and other aspects of cognition, such as memory, that tend to fluctuate in presence and severity. Patients may exhibit signs and symptoms of anxiety or psychosis, including delusions, hallucinations, and altered mood. They often have altered arousal and psychomotor activity, ranging from reduced responsiveness at near-coma level to hyper-vigilance, and agitation. Patients with delirium show characteristic EEG changes, in particular slow wave activity, and a disrupted sleep-wake cycle^25^. Mechanistic understanding of delirium in humans in general and in AKI is very limited, and acute delirium after AKI has not been studied previously in pre-clinical models. We therefore established a model of AKI-induced acute delirium using bilateral renal ischemia reperfusion-induced tissue injury as a model tissue injury with uremia. We used young adult (10-12 week old) C57Bl6 *wt* mice throughout our studies. We performed sham or AKI surgeries followed by behavioral testing, as well as flow cytometry and histology of cortex and hippocampus (**Figure 1A**). Serum blood-urea-nitrogen (BUN) levels were appropriately elevated at 48 hours after AKI as compared to sham animals, indicating significant kidney injury and uremia (**Figure 1B**). The elevated plus maze (EPM) is a commonly preferred test to assess for anxiety-like behavior, risk aversion in mice, and measures the conflict between the innate drive to explore a novel environment and the fear of open, elevated spaces. It also tests locomotion. In EPM testing, AKI animals showed significantly reduced exploration, time spent in open arms, and total distance travelled (**Figure 1C**). The reduction in locomotion in AKI animals is unlikely the result of post-operative pain as sham animals underwent the same surgical procedure except for clamping of the renal vessels and also received the same post-surgical analgesia. The open field (OFT) test evaluates locomotion (distance, speed) and anxiety based on the animal’s preference to stay near the walls (thigmotaxis) versus exploring the bright, exposed center. While thigmotaxis was not different, locomotion (distance, speed) was also reduced in AKI compared to sham animals in the OFT test (**Supplemental Figure 1A**). The abbreviated fear conditioning (FC) test assesses several neurological capacities: Associative learning (Day 1), by testing the cognitive ability to link a neutral stimulus (tone or light) with an aversive event (mild foot shock). Contextual memory (Day 2), by evaluating hippocampus-dependent memory when the animal recognizes the physical chamber where the training occurred. Fear expression (Day 1+2), by quantifying defense responses, specifically measured through the duration of freezing behavior. Mice with AKI had increased freezing during both the conditioning (Day 1) and contextual recall testing (Day 2) as compared to sham animals, also consistent with anxiety-like behavior(**Figure 1D**). Taken together these behavioral findings support the existence of an anxiety-like state in AKI mice, possibly linked to hippocampal inflammation, and combined with a reduction in locomotion that could be interpreted as reduced psychomotor regulation, both features of delirium that can be observed in humans^25^. To further assess global impact of AKI on brain activity, we performed EEGs in sham and AKI mice. EEG testing revealed that AKI, but not sham mice exhibited disturbance in sleep-wake cycle with increased NREM sleep during the dark phase, decreased wakefulness, and increased wake-NREM transitions. Sleep-wake cycle disturbance detected by EEG is also seen in human delirium^25^ and in sepsis-induced delirium in rodents^26^. AKI animals showed an increase in Theta power in the dark and no changes in Delta power (**Figure 1E, Supplemental Figure 1B**). Humans with delirium, in contrast, exhibit increases in Theta and Delta power^25,27^. The interpretation of our EEG findings in mice and comparison with human EEG is not straightforward, as in mice Theta waves are related to faster behavioral or functional states. In surgery- or sepsis-associated encephalopathy (delirium) rodents show decreased Theta and increased Delta power^28,29^, opposite to our model. Since AKI-induced EEG changes have not been studied in rodents previously, the significance of this discrepancy is at least unclear. Yet, our model provides evidence of changes in cortical function, in part similar to delirium in humans. Taken together, these results validate the translational relevance of our pre-clinical model of AKI-induced delirium in mice.

**Fig. 1.**
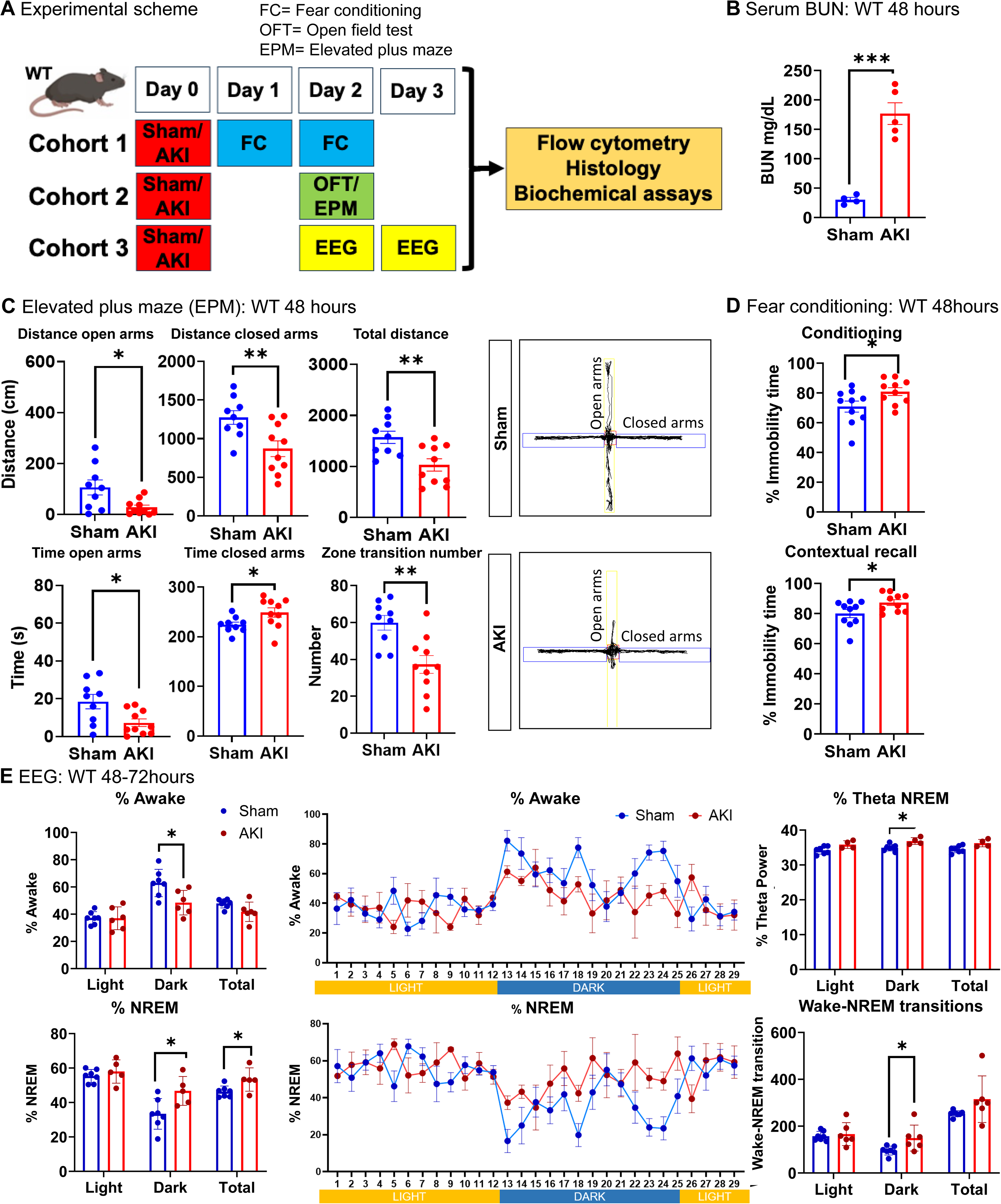
AKI induces acute neurological dysfunction with delirium-like symptoms in mice. (**A**) Experimental scheme: Behavioral testing and EEG analysis were performed in young adult (10-12 weeks) wt mice between 24-72 h after AKI as indicated followed by sacrifice and tissue harvesting for flow cytometry, histology and biochemical analysis. (**B**) Serum BUN values 48h hours after sham or AKI. (**C**) Elevated plus maze test 48 hours after AKI. (**D**) Contextual fear conditioning test done at 24 hours (Day 1 conditioning) and 48 hours (Day 2 contextual recall) after sham/AKI. (**E**) EEG recording at 48-72 hours after sham/AKI. (*n* = 7 to 10 animals per measurement. \**P* < 0.05, \*\**P* < 0.01, \*\*\**P* < 0.001 by student t-test. All data represent mean ± SEM.

### AKI induces acute hippocampal monocytic neuroinflammation without significant changes in blood brain barrier permeability

We next examined the hypothesis that AKI induces neuroinflammation. Following behavioral testing mice were sacrificed and perfused with PBS before brains were isolated and subjected to flow cytometric analysis of immune cells and histology. Single-cell preparations of the hippocampus and cortex at 48 hours after AKI compared to sham did not show a difference in overall CD45^+^ immune cell counts but showed a significant increase of CD11b^+^Ly6C^+^ monocytes in the hippocampus, but not in the cortex. The AKI-induced hippocampal monocyte accumulation consisted of Ly6C^high^ monocytes known to represent CCR2^+^ monocytes recruited from the circulation, as well as Ly6C^low^ monocytes (**Figure 2A**). The latter may at least in part represent formerly CCR2^+^ monocytes that are in the process of developing into macrophage- or microglia-like cells. To test this hypothesis, we employed CCR2 fate mapping, using CCR2-ERT2Cre-Rosa26tdTomato^flox/flox^ mice, in which tamoxifen administration irreversibly activates tdTomato expression in CCR2^+^Ly6C^high^ cells and their progeny. One dose of tamoxifen in CCR2-ERT2Cre-Rosa26tdTomato^flox/flox^ mice marked about 60% of circulating CCR2^+^ Ly6C^high^ monocytes for 3-5 days (**Supplemental Figure 2A**). In tamoxifen-treated AKI mice roughly 70% of Ly6C^high^ circulating monocytes (**Figure 2B**) and 11% of Ly6C^high^ hippocampal monocytes were tdTomato^+^, compared to about 50% and 4% respectively in sham mice. Some tdTomato^+^ cells in the hippocampus were Ly6C^low^ or even Ly6C^-^ (**Figure 2C**). Some hippocampal tdTomato^+^ cells stained partially positive for the microglia marker ionized calcium-binding adaptor molecule 1 (Iba1), but not for the neuronal nuclear antigen (NeuN) (**Figure 2D**). This suggests that AKI-induced hippocampal CCR2^+^ monocytes develop into microglia-like progeny.

**Fig. 2.**
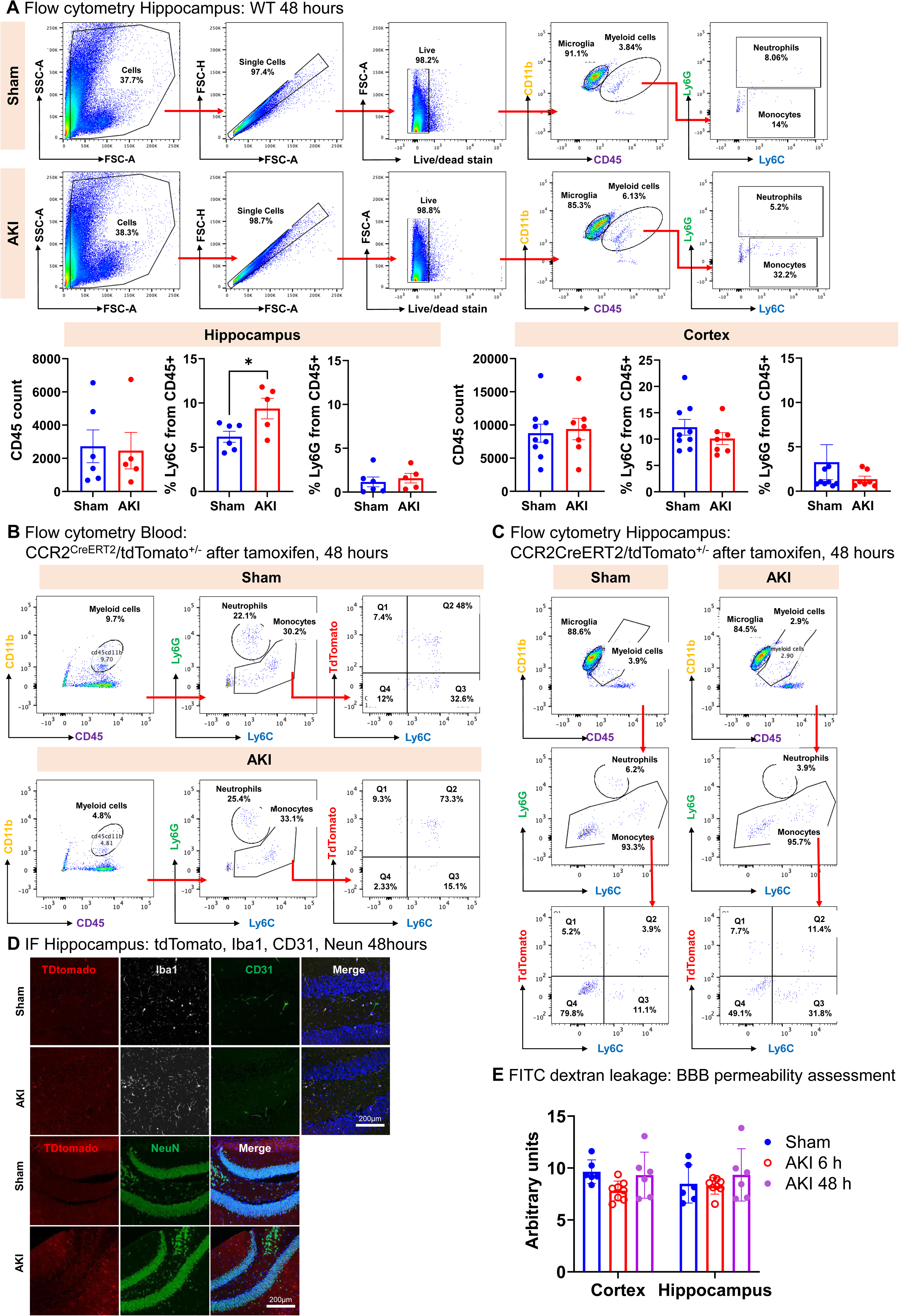
AKI induces acute hippocampal monocytic neuroinflammation without significant changes in blood brain barrier permeability. (**A**) Hippocampus and cortex flow cytometry analyses at 48 hours after sham/AKI surgery in young adult *wt* mice. Representative gating strategy (upper panel), total immune cell counts, percentage of monocytes and neutrophils respectively for hippocampus (on the left) and cortex (on the right). (**B**) Flow cytometry blood analysis of CCR2^CreERT2tdTomato+/-^ mice injected with tamoxifen and examined at 48hours after sham/AKI surgery. Shown is the gating strategy for the detection of tdTomato^+^ cells in the circulation. (**C**) Flow cytometry assay of hippocampus of CCR2^CreERT2tdTomato+/-^ mice done at 48hours after sham/AKI surgery. Gating strategy to detect tdTomato^+^ LyC^hi^ and Ly6^low/negative^ cell populations. (**D**) Immunofluorescence staining of hippocampal tissue from CCR2^CreERT2tdTomato+/-^ mice collected at 48hours after sham/AKI. Staining shows that tdTomato^+^ cells partially colocalize with Iba1; a marker for microglial cells, but not with Neun; a neuronal marker, or CD31, an endothelial marker. (**E**) Blood brain barrier permeability assessment at 6 hours and 48 hours after sham/AKI surgery in *wt* mice in the hippocampus and cortex regions. (*n* = 6 to 10 animals per measurement. \**P* < 0.05, \*\**P* < 0.01, \*\*\**P* < 0.001 by student t-test. All data represent the mean ± SEM.

How these cells may be connected to neurologic dysfunction after AKI needs further study. We did not observe changes in microglial activation via flow cytometry analysis or 3D morphologic reconstructions of Iba^+^ cells in the hippocampus at this early (48 hours) time point after AKI (**Supplemental Figure 2B**). As increases in blood brain barrier (BBB) permeability have been proposed to play a role in AKI-induced neuroinflammation^18,25^ and could permit increased access of uremic toxins to the brain, but also an increase in monocyte infiltration, we tested whether AKI induced BBB permeability changes occur in our model. Using intravenous injection of 70kd FITC-Dextran, we did not observe blood brain barrier permeability changes at 6 or 48 hours after AKI as compared to sham animals (**Figure 2E**). These findings suggest that the observed neuroinflammation and acute neurologic dysfunction occur independently of significant changes in BBB permeability.

### Circulating osteopontin drives acute hippocampal monocytic neuroinflammation

We previously reported that circulating osteopontin (OPN/SPP1) released by the injured kidney after AKI causes remote lung inflammation by acting on lung intravascular CCR2^+^ monocytes that develop into lung interstitial macrophage progeny^23,24^, a phenotype reminiscent of the CCR2^+^ monocyte-driven microglia-like cell accumulation in the hippocampus we observed after AKI. Uremia alone (bilateral nephrectomy) did not induce this kidney-lung remote inflammation phenotype, and anti-OPN antibody or global OPN knockout (OPN-KO) prevented it^23^. We thus tested whether OPN could also be involved in our remote hippocampal inflammation phenotype in the kidney-brain axis. In AKI mice compared to sham, OPN mRNA and protein levels were elevated in the kidney (**Figure 3A**) and serum (**Figure 3B**) but OPN mRNA or protein were not elevated in the hippocampus or cortex (**Figure 3C+D**). This is similar to our kidney-lung phenotype where OPN is upregulated in kidney and serum, but not in the lung, and kidney transplantation experiments conclusively showed that during AKI the kidney represents the sole source of circulating OPN^23^. OPN-KO animals with AKI compared to sham controls, or AKI wt animals treated with OPN neutralizing antibody compared to IgG control showed AKI with significant serum BUN elevations, at levels very similar to *wt* animals (**Figure 3E+F** compare to **Figure 1B**). However, despite biochemical evidence of uremia, neither hippocampal monocyte accumulation (**Figure 3G+H**) nor behavioral abnormalities in EPM were detected (**Figure 3I+J**). We tried to raise serum OPN levels in OPN-KO mice by either injecting it, by using osmotic minipumps, or by using adenovirus-driven OPN overexpression under a liver-specific promotor in an attempt to revert their protection and to test whether OPN alone without AKI and uremia can induce neuroinflammation and delirium-like symptoms. However, due to the known very short half-life of OPN, even when pegylation was added to extend half-life, we were unable to reach significant serum levels. AAV8-driven OPN overexpression under the control of a liver-specific promotor induced liver failure due to failure to repair the viral infection (**Supplemental Figure 3**). OPN indeed has known functions in tissue repair^30^. Still, taken together, our results suggest that AKI-induced CCR2^+^ monocyte-driven hippocampal inflammation and delirium-like symptoms depend on circulating OPN, but not primarily on uremia, in both instances similar to remote lung inflammation.

**Fig. 3.**
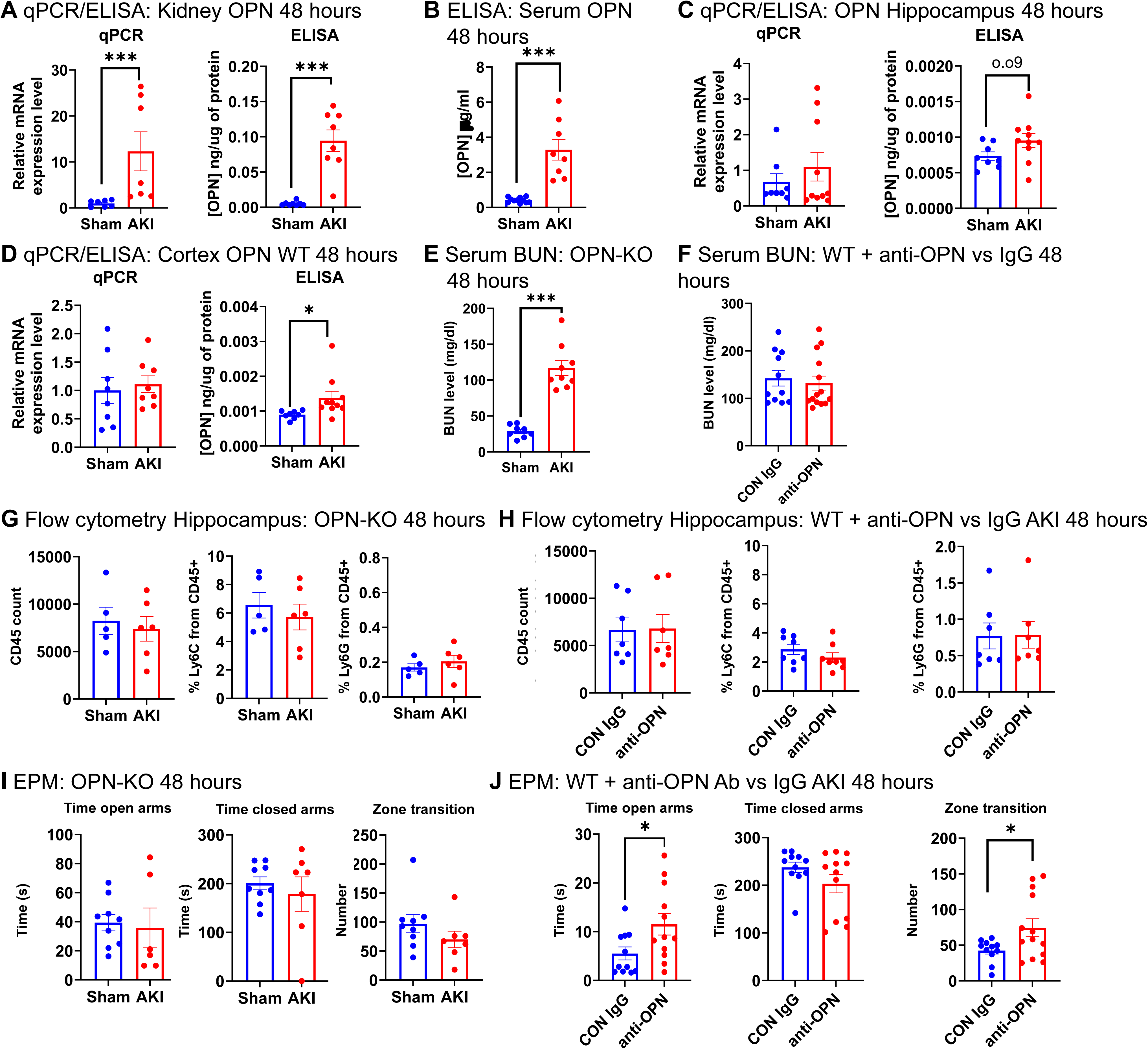
Circulating osteopontin drives acute hippocampal monocytic neuroinflammation. (**A**) Osteopontin mRNA (qPCR) and protein expression (ELISA) in the kidney at 48 hours after sham/AKI in *wt* mice. (**B**) Serum osteopontin (OPN) ELISA in *wt* mice. (**C**) Osteopontin mRNA (qPCR) and protein expression (ELISA) in the hippocampus at 48hours after sham/AKI in *wt* mice. (**D**) Osteopontin mRNA (qPCR) and protein expression (ELISA) in the cortex at 48hours after sham/AKI in *wt* mice. (**E**) Serum BUN levels in OPN-KO mice at 48 hours after sham/AKI. (**F**) Serum BUN levels in *wt* AKI mice injected with either control IgG or anti-OPN antibody at 48 hours after sham/AKI. (**G**) Hippocampal flow cytometry at 48hours after sham/AKI in OPN-KO mice. (**H**) Hippocampal flow cytometry at 48 hours after AKI in *wt* mice injected with control or anti-OPN antibody. (**I**) Elevated plus maze test of OPN-KO mice at 48 hours after sham/AKI. (**J**) Elevated plus maze test of *wt* AKI mice injected with control or anti-OPN antibody at 48 hours after AKI. (*n* = 6 to 10 animals per measurement. \**P* < 0.05, \*\*\**P* < 0.001 by student t-test. All data represent the mean ± SEM.

### Circulating OPN fragments generated by thrombin cleavage in the lung drive AKI-induced neuroinflammation and delirium

OPN-R153A (OPN-KI) mice carry a point mutation in OPN that make it resistant to thrombin cleavage, preventing production of its two potential signaling fragments, OPN-N and OPN-C, also detectable in human biological fluids^31^. In western blots of serum samples from AKI *wt* mice compared to sham controls we detected two cleavage products of OPN at the predicted molecular size for OPN-N and OPN-C; these OPN fragments were not detected in OPN-KI mice, the latter suggesting that thrombin cleavage of circulating OPN may be required for AKI-induced remote neuroinflammation (**Figure 4A**). Consistently, although serum BUN elevations were similar to *wt* littermates and OPN-KI mice with AKI (**Figure 4B** compare to **Figure 1B**), AKI-induced hippocampal monocytic infiltration (**Figure 4C**) and anxiety-like behavior in EPM (**Figure 4D**) did not occur in OPN-KI mice (**Figure 4E**), compared to *wt* animals (compare to **Figure 1C+2A**), implicating circulating thrombin cleaved OPN fragments. We performed specific thrombin activity assays to determine where in the body OPN cleavage by thrombin may occur. We had observed significant lung capillary thrombosis in our AKI-induced remote lung inflammation phenotype^23^ and suspected the lung as the site of thrombin activation and OPN cleavage. We further considered that such thrombotic complications would likely be very risky in the brain vasculature; we did however not observe any gross thrombotic complications in the brain in our experiments (not shown). Using a specific thrombin activity assay, we examined thrombin activity in protein lysates from perfused organs and serum samples isolated from sham and AKI *wt* mice. Thrombin activity in serum was highest overall but was moderately decreased by AKI. AKI-induced increased thrombin activity was only detected in the lung, but not the brain, kidney, liver, or spleen. (**Figure 4F**). The measured thrombin activity in the lung was completely inhibited with addition of the highly specific naturally occurring thrombin inhibitor hirudin (**Figure 4G**), showing high specificity of the assay for thrombin cleavage. In summary, our data suggests that cleavage of kidney-released circulating OPN by thrombin in the lung vasculature is required to generate OPN fragments that drive hippocampal neuroinflammation and delirium-like symptoms in uremic mice.

**Fig. 4.**
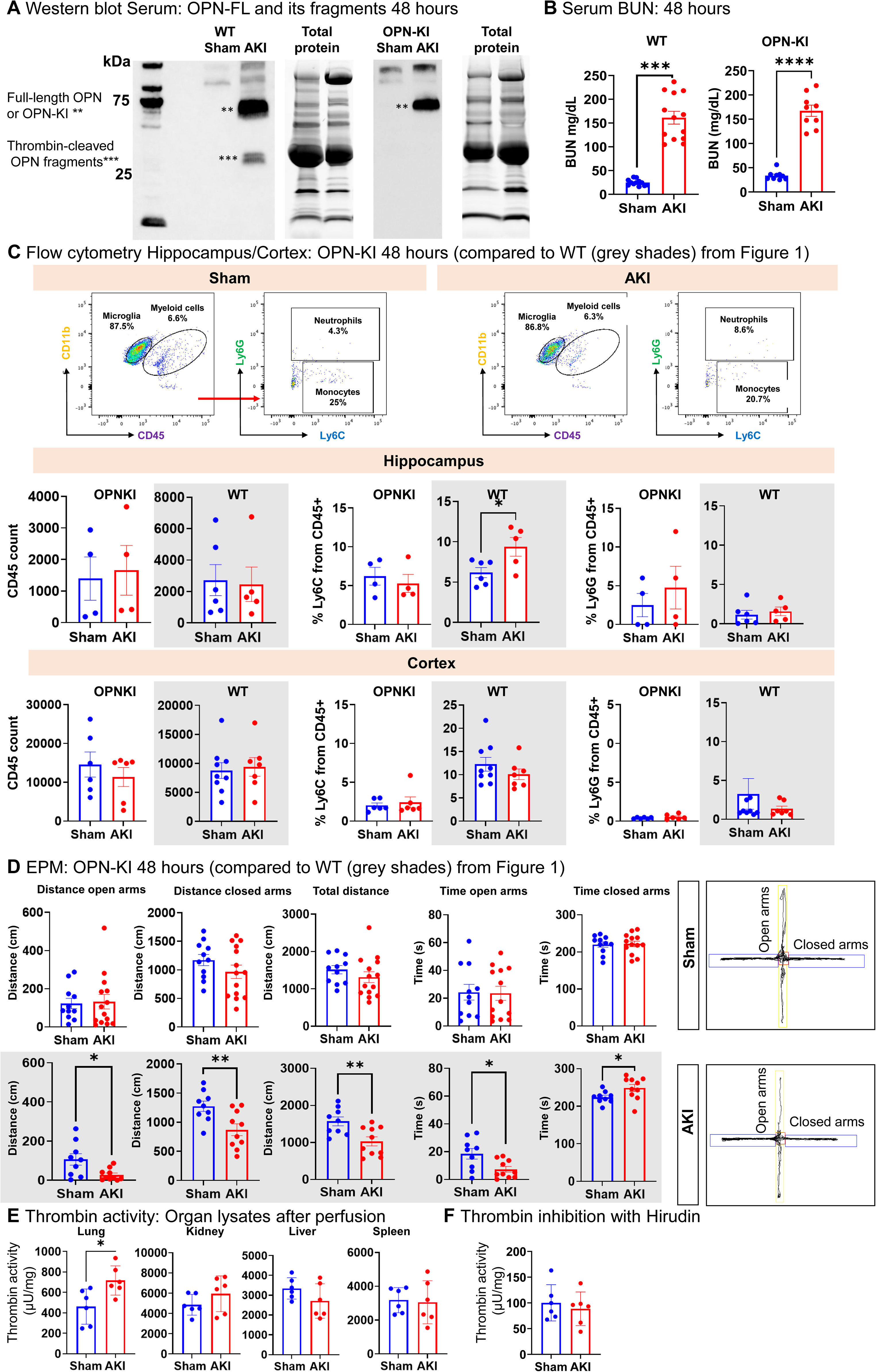
Thrombin cleavage of circulating OPN drives AKI-induced neuroinflammation and delirium. (**A**) Western blot of serum from *wt* and OPN-KI sham/AKI mice. Both *wt* and OPN-KI AKI mice show significant increase of full-length OPN compared to sham serum, but only wt mice show thrombin-cleaved OPN fragments. (**B**) Serum BUN levels in *wt* and OPN-KI mice at 48 hours after sham/AKI. (**C**) Hippocampal and cortical flow cytometry analyses at 48 hours after sham/AKI in OPN-KI mice. (**D**) Elevated plus maze test of OPN-KI mice at 48 hours after sham/AKI. Shaded graphs repeat *wt* data from Figure 1 for direct comparison (*n* = 6 to 10 animals per measurement. \*\*\**P* < 0.001 by student t-test. All data represent the mean ± SEM.

### Uremic retention solutes are equally elevated in mice with or without neuroinflammation and delirium

Since we observed that *wt* and OPN-KO or OPN-KI animals showed the same degree of uremia measured by serum BUN elevation after AKI (**Figure 5A** compare to **Figures 1B, 3B, 4C**), but only wt animals showed remote hippocampal inflammation and delirium-like symptoms, we further investigated the role of uremic toxins in our model. We determined serum uremic retention solute levels broadly by mass spectrometry in sham/AKI *wt*, OPN-KO, and OPN-KI mice, including a number of solutes correlatively linked to chronic neurocognitive dysfunction in CKD, in particular indoxyl and p-cresol sulfate^32,33^. Uremic toxins were equally elevated in AKI *wt*, OPN-KO, or OPN-KI mice, including uremic toxins said to cause neurologic dysfunction^34–36^ (**red font in heatmap in Figure 5A**), in particular indoxyl and p-cresol sulfate (**Figure 5B**). However, Hippurate was not even upregulated in the AKI groups, questioning its’ role in neurological dysfunction. Our pre-clinical model thus challenges the notion that uremic retention solutes are major drivers of acute AKI-induced neuroinflammation and delirium. If they played a role, it requires OPN.

**Fig. 5.**
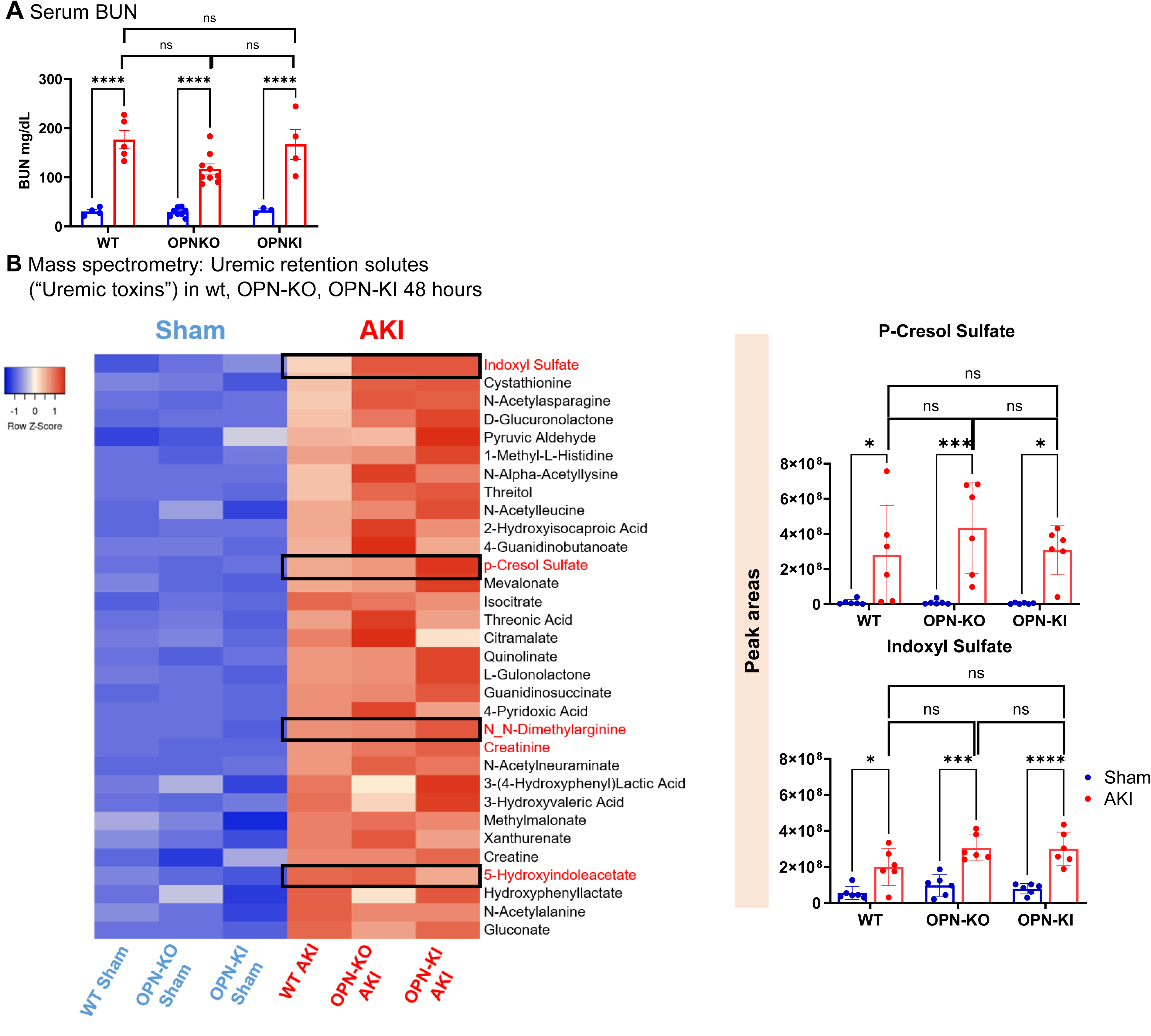
Uremic retention solutes are equally elevated in mice with or without neuroinflammation and delirium. (**A**) Serum BUN levels in *wt,* OPN-KO, and OPN-KI at 48hours after sham/AKI mice. **B**) Uremic retention solutes were measured by mass spectrometry. (*n* = 5 to 6 animals per measurement. \**P* < 0.05, \*\**P* < 0.01, \*\*\**P* < 0.001 by two-way ANOVA followed by Šídák’s multiple comparisons test. All data represent the mean ± SEM.

### OPN fragments drive monocytic hippocampal inflammation and delirium-like symptoms via induction of chemokine-ligand-2 (CCL2) in the hippocampus

To investigate the OPN-induced signal that drives hippocampal monocyte accumulation, we tested for AKI-induced upregulation of monocyte chemoattractants in the hippocampus. In AKI *wt* mice compared to sham, monocyte chemo-attractant chemokine-ligand-2 CCL2 mRNA (also called monocyte-chemoattractant-protein-1, MCP-1), a key ligand for CCR2 receptor and a tissue recruitment signal for CCR2^+^ monocytes, was robustly upregulated in the hippocampus by 48 hours after AKI, as detected by qPCR, with a trend to increase in the cortex (**Figure 6A** and **Supplemental Figure 4**). Consistent with their lack of hippocampal monocyte accumulation and delirium-like symptoms, OPN-KO or OPN-KI mice did not show upregulation of CCL2 mRNA in the hippocampus or cortex (**Figure 6B**). Hippocampal CCL2 mRNA at increased levels compared to sham control could also be detected by in-situ hybridization (CCL2 RNAscope) (**Figure 6C**). In CCL2-RFP^flox/flox^ mice CCL2 expression can be detected by co-expression of red fluorescent protein (RFP) in any CCL2 expressing cell. In these mice an aphthovirus 2A “cleavage site” is inserted between CCL2 and RFP. This short 16–18 amino acid peptide sequence from the foot-and-mouth disease virus (FMDV) triggers a co-translational ribosomal “peptide-bond-skipping” (StopGo) event rather than true proteolytic cleavage, separating upstream and downstream proteins from a single open reading frame. Thus, CCL2 is separated from RFP and CCL2 function and biology remain unaffected^37^. Using CCL2-RFP mice, we detected AKI-induced upregulation of CCL2 exclusively in CD31^+^ endothelial cells in the hippocampus and cortex (**Figure 6D** and **Supplemental Figure 4B**). We then tested whether CCL2 is required to attract monocytes to the hippocampus after AKI. Anti-CCL2 neutralizing antibody compared to control IgG did not affect kidney injury **(Figure 6E)** but prevented hippocampal monocyte accumulation (**Figure 6F**) and delirium-like symptoms based on EPM findings (**Figure 6G**), validating CCL2’s critical role in AKI-induced remote neuroinflammation.

**Fig. 6.**
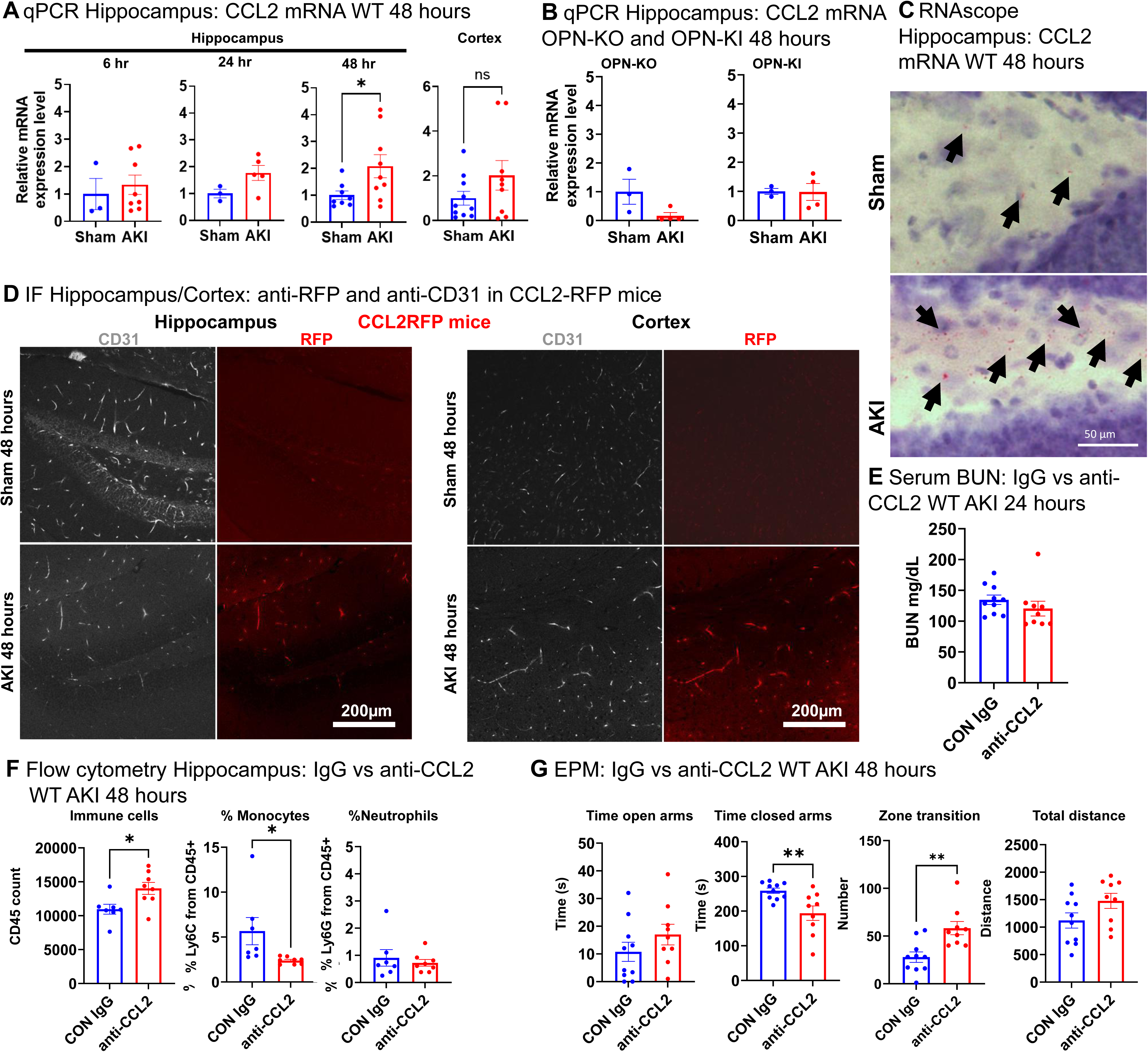
OPN fragments drive monocytic hippocampal inflammation and delirium-like symptoms via induction of chemokine-ligand-2 (CCL2) in the hippocampus. (**A**) Hippocampal and cortical CCL2 mRNA expression (qPCR) at 6, 24, and 48 hours (hippocampus) and at 48 hours (cortex) after sham/AKI surgery in *wt* mice. (**B**) Hippocampal CCL2 mRNA expression (qPCR) at 48 hours after sham/AKI in OPN-KO and OPN-KI mice. (**C**) Hippocampal CCL2 mRNA *wt* sham/AKI mice assessed with in-situ hybridization (RNAscope). (**D**) Immunofluorescence staining showing RFP signal (indicating CCL2 protein expression) colocalizes with CD31 (marker for endothelial cells) in the hippocampus and cortex of CCL2-RFP^flox/flox^ AKI mice. (**E**) Serum BUN level at 48 hours after AKI in *wt* AKI mice injected with control IgG or anti-CCL2 antibody. (**F**) Hippocampal flow cytometry analysis in *wt* AKI mice injected with control IgG or anti-CCL2 antibody at 48 hours after AKI. (**G**) Elevated plus maze analysis in *wt* AKI mice injected with control IgG or anti-CCL2 antibody at 48 hours after AKI. (*n* = 4 to 10 animals per measurement. \**P* < 0.05, \*\**P* < 0.01, \*\*\**P* < 0.001 by student t-test. All data represent the mean ± SEM.

### Both thrombin-cleaved OPN fragments together induce chemokine-ligand-2 (CCL2) in hippocampal CCR2^+^ monocytes and their interstitial macrophage-like progeny

Earlier (**Figure 2**) we had fate-mapped CCR2^+^ monocytes in our model using tdTomato fluorescence (CCR2-ERT2Cre-Rosa26tdTomato^flox/flox^) and discovered that AKI-induced tdTomato^+^ cells in the hippocampus consist of CCR2^+^Ly6C^high^ monocytes, but also tdTomato^+^Ly6C^intermeditate/low^ monocytes and tdTomato^+^Ly6C^negative^ cells, the former representing circulating tdTomato^+^CCR2^+^ monocytes recruited to the hippocampus and the latter their progeny. Based on our published kidney-lung experiments, CCR2^+^ monocytes are very likely target cells of OPN^23,38^. However, based on our CCL2 experiments endothelial cells also represent candidate target cells. We thus hypothesized that CCR2^+^ monocytes and/or their hippocampal progeny, possibly microglia-like cells (**Figure 2 D**), may be target-cells of OPN fragments. We thus tested the ability of OPN fragments to induce CCL2 in interstitial macrophage-like cells or in primary lung endothelial cells. Thioglycolate-elicited peritoneal macrophages are derived from circulating CCR2^+^ monocytes and can serve as a model for macrophage-like cells in the hippocampus. We generated mouse OPN expression vectors encoding OPN-FL, OPN-N, OPN-C, and thrombin uncleavable OPN-R153A (OPN-KI), including GST-fusion and epitope tags as indicated (**Figure 7A**). We expressed them in HEK293T cells and purified them from cell lysate using glutathione sepharose beads (**Figure 7B**). We removed GST by cleavage with thrombin or Factor Xa in the case of OPN-FL and detected them by anti-epitope tag WB; OPN-N is detected as two isoforms (**Figure 7C**). Only OPN-N+C but not OPN-FL, OPN fragments alone, or OPN-KI protein, effectively induced CCL2 in CCR2^+^ monocyte-derived peritoneal macrophages on the mRNA and protein level at 24 hours (**Figure 7D**), or on the mRNA level in primary mouse endothelial cells (**Figure 7E**). The same effect was not observed in a microglial cell line (**Supplemental Figure 5**). Taken together, this data supports the hypothesis that OPN fragments act on receptors in circulating or hippocampal CCR2^+^ monocytes and their microglia-like progeny and/or on endothelial cells to induce CCL2. It is also possible that OPN fragment-activated CCR2^+^ monocytes drive endothelial cells to produce CCL2 by yet to be determined signals.

**Fig. 7.**
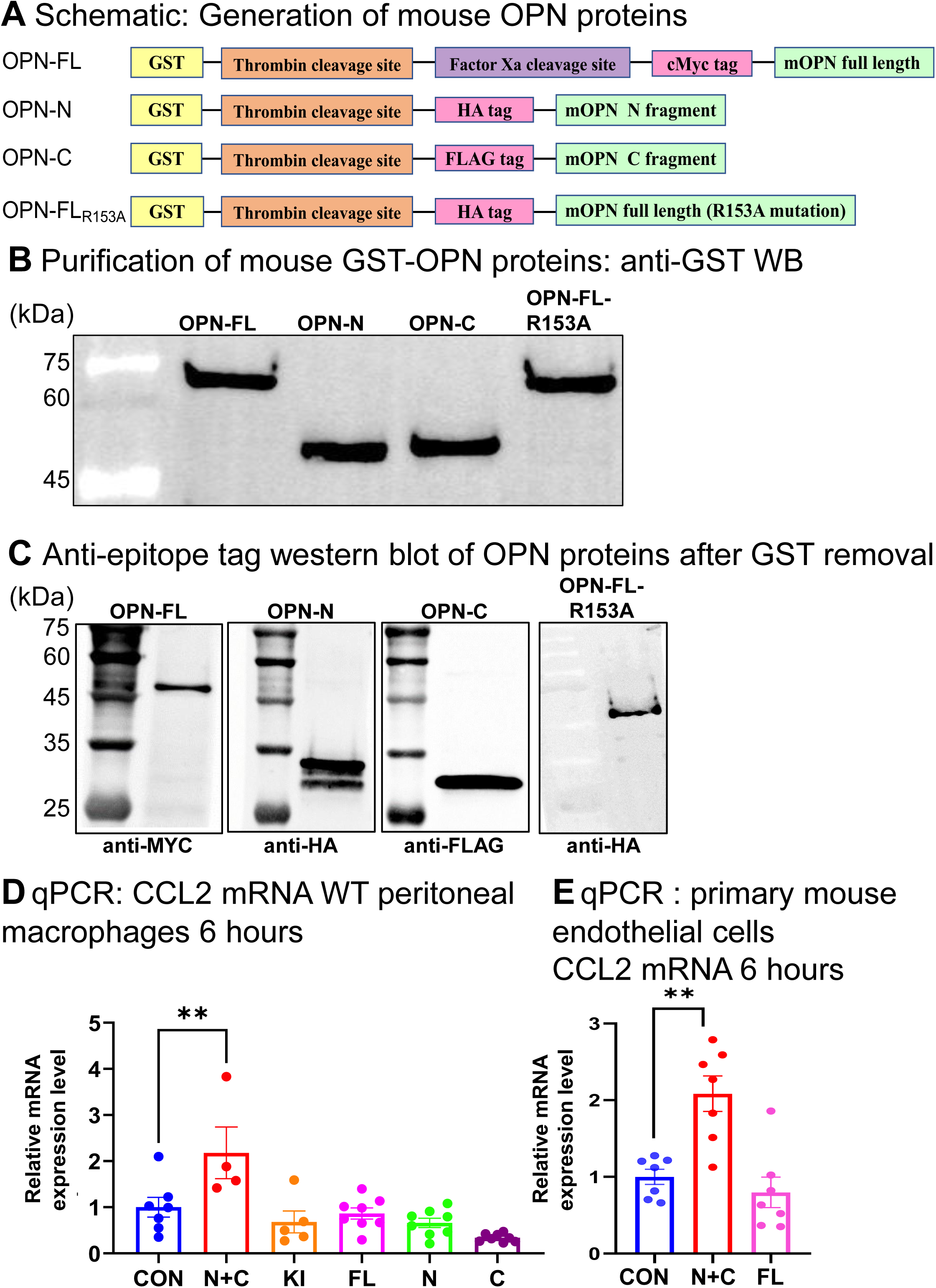
OPN fragments drive monocytic hippocampal inflammation and delirium-like symptoms via induction of chemokine-ligand-2 (CCL2) in the hippocampus. (**A**) Mouse OPN constructs. Note factor Xa cleavage site in OPN-FL to remove GST; thrombin used otherwise. (**B**) anti-GST western blot of OPN-FL, OPN-N + OPN-C fragments and thrombin uncleavable OPN-R153A. (**C**) anti-epitope tag western blot of OPN proteins. (**D**) CCL2 mRNA expression in peritoneal macrophages after 6 hours of treatment with different OPN full length and thrombin-cleaved OPN fragments. Only N+C fragment treatment produced significant level of CCL2 expression (qPCR). (**E**) CCL2 mRNA expression (qPCR) in mouse primary endothelial cells after 6 hours of treatment with different OPN full length and OPN N+C fragments. Again, only N+C fragment treatment produced significant level of CCL2 expression (qPCR). **p<0.01, (*n* = 4 to 7 samples per measurement.). One way ANOVA followed by post hoc Tukey. All data represent the mean ± SEM.

## Discussion

Alterations of mental status attributed to acute uremia (uremic toxins) have been described since 1839, when Addison reported dullness, sluggishness, drowsiness, stupor and coma, but also convulsions and psychotic manifestations in patients with acute renal failure^39^. Locke published a detailed modern characterization of AKI-induced delirium and neurological abnormalities in 1961, studying 13 patients with mostly severe AKI of various causes and no prior history of kidney disease. Almost 40% required dialysis, attesting to disease severity in this small study. All patients had delirium and some degree of muscle weakness, and variably exhibited lethargy, stupor or coma, or anxiety, agitation, irritability and confusion. EEGs showed “coma”-like activity with slow waves, alpha rhythm dropout and excess theta and delta power which normalized when patients recovered from delirium; some patients experienced seizures. Disturbance of sleep-wake cycle was common^40^. Similar EEG observations were made by Klinger et al. in 23 patients with uremia and delirium^41^. Our mouse model of AKI-induced delirium corroborates these human findings: we also observed delirium-like symptoms, reduced locomotor activity, and EEG changes characteristic of delirium, highlighting the translational validity of our model. How AKI acutely impairs neurologic function in humans or animals is largely unknown, impeding development of therapies to mitigate delirium risk after AKI. Several mechanisms have been proposed, including uremic toxins accumulating due to reduced renal clearance and upregulation of systemic cytokine-mediated inflammatory processes linked to neuroinflammation, both possibly enhanced by BBB disruption^18^.

In those early human AKI studies, blood chemistries, including serum electrolytes, bicarbonate, pH, creatinine and BUN (a surrogate for uremic toxin levels), correlated with neither delirium nor delirium-linked EEG abnormalities^40,41^, questioning a direct mechanistic link between uremic toxins and acute AKI-induced delirium. Indeed, this hypothesis rests very largely on correlative data from chronic kidney disease (CKD) patients or pre-clinical CKD models, rather than on studies in AKI patients or pre-clinical AKI models^18,36^. How uremic toxins cause delirium is unclear; hypotheses include direct neurotoxicity and/or induction of neuroinflammation^36,42^. One study reported that dialysis reduced delirium risk in severe but not milder AKI, lending limited support to involvement of a dialyzable factor^43^. Uremic toxins are a highly diverse group of at least 150 molecules^34,36,44^. Metabolic profiling correlated four uremic metabolites (4-phenylacetylglutamine, phenylacetylglutamine, hippurate and prolyl-hydroxyproline) with chronically impaired executive functioning in dialysis-dependent CKD patients^45^. Although these substances are in principle easily dialyzable, except for hippurate, which is moderately protein bound, dialysis struggles to remove them fully owing to their high generation rate and its inability to match the excretion rate of a functioning kidney. Indoxyl sulfate and p-cresyl sulfate, the two most studied gut bacteria-derived uremic toxins linked to chronic neurocognitive dysfunction in dialysis patients and pre-clinical models, are highly protein bound and not easily dialyzable^36,46–51^. Whether they are linked to acute AKI-induced delirium is unknown. Whether dialysis improves acute delirium in AKI patients remains uncertain based on retrospective and observational data and has not been studied directly^18^. Reduced consciousness, asterixis, and acute delirium are typically slow to reverse, beginning 24-48 hours after starting dialysis, often improving over several days to weeks and lagging behind serum BUN improvement^18^. This is hypothesized, though unproven, to reflect slow or absent clearance of middle-sized (>20–25kDa) and protein-bound uremic toxins, which are particularly hard to remove with commonly used dialysis technology. Human OPN and its fragments are also in this size range (25-35kDa; full-length OPN is around 70kDa)^52,53^ and would be difficult to remove with standard hemodialysis. OPN has not been studied in hemodialysis effluent but has been found in peritoneal dialysis effluent^54^. However, peritoneal dialysis is infrequently used for AKI in the western world, and no data on its effectiveness in delirium are available. In our model, uremic toxins elevated by AKI, including those linked to neurocognitive dysfunction above, were equally elevated in AKI wt, OPN-KO or OPN-KI animals, yet only AKI wt animals showed delirium-like symptoms. Thus, based on our pre-clinical model and the available clinical data, uremic toxins may not have the prominent mechanistic role previously ascribed to them in acute AKI-induced delirium. They may, however, still be required for the effect of OPN. We could not resolve this, as serum OPN elevation could not be separated experimentally from AKI and uremic toxin elevation. Our results do not exclude a role for uremic toxins in chronic neurocognitive dysfunction in AKI survivors or CKD patients.

The hypothesis that neuroinflammation is central to delirium rests on several observations^55–57^. Systemic inflammation is a feature of many medical or surgical conditions associated with delirium that involve sterile tissue injury (e.g. AKI, myocardial infarction, surgical or non-surgical trauma) and/or injury from infection^56^. Infection typically starts locally (e.g. pneumonia, skin abscess) but can disseminate systemically (sepsis). Sterile and infection-related tissue injury produce similar or even identical local inflammatory mediators and cytokines. Early on tissue injury and infection drive release of alarmins, such as pathogen- and danger-associated molecular patterns (PAMPS, DAMPS), that activate the innate immune system via Toll-like receptors, and of classical cytokines such as interleukin-1β (IL-1β), tumor necrosis factor (TNF), and interleukin-6 (IL-6)^55,58,59^. These molecules relay peripheral inflammation to the central nervous system (CNS) and induce a CNS inflammatory response (neuroinflammation) via at least two mechanisms that likely act together: activation of local afferent trigeminal or vagal pathways through neuro-immune interactions, which signal peripheral tissue injury and infection to the CNS; and entry into the circulation, allowing inflammatory mediators to reach the brain^55,60–62^, as studied here for OPN. Cytokine uptake into the brain from the circulation occurs by several mechanisms: entry at the choroid plexus and circumventricular organs that lie outside the BBB; active endothelial transport and uptake at the BBB, as shown for IL-1β, TNF, or IL-6 and others^63–65^, and/or by enhanced BBB leakiness for example in the setting of sepsis^55,66^. CNS resident cells that associate with the neuro-vascular barrier unit^67^ respond to these stimuli with inflammatory activation and/or proliferation: endothelial cells, pericytes, astrocytes, perivascular macrophage-like cells and microglia^57^. This propagates systemic inflammatory stimuli into the brain, influencing synaptic transmission, neuronal excitability, and cerebral blood flow^68^. Together, these coordinate a complex adaptive CNS response collectively referred to as “sickness behavior” that aids organismal defense against acute peripheral tissue injury and/or infection^55–57,69^. Delirium represents a maladaptive consequence of this adaptive response. While much remains unknown, several studies suggest that core symptoms of delirium, in particular impaired consciousness, involve changes in neuronal activity and brain network connectivity that impair information integration by functionally disconnecting brain regions^70,71^. Further studies are needed to clarify the exact neuropathological mechanism(s) by which AKI and OPN induce disturbance of brain networks and delirium.

There is limited direct evidence for neuroinflammation in human delirium. In a post-mortem case-control study of 9 patients with delirium and 6 age-matched controls, those with delirium showed significant increases in markers of microglial (HLA-DR, CD68) and astrocyte (GFAP) activity, and in IL-6 immunoreactivity across all brain areas^72^. Another post-mortem study correlated systemic inflammation with endothelial and perivascular cell activation in human brain tissue^73^. Rodent studies using lipopolysaccharide (LPS), cytokines and surgical procedures established that peripheral immune stimulation does induce central neuroinflammation with acute, transient “sickness behavior” characterized by sleepiness, anorexia, apathy and cognitive impairment^69^. Bacterial sepsis or systemic injection of the PAMP LPS, a major outer membrane component of Gram-negative bacteria, can induce hippocampal and cortical inflammation with microglial response, local cytokine upregulation, and delirium-like symptoms in mice^56,57,74,75^, including EEG findings characteristic of delirium^76,77^, very similar to the observations we made with sterile tissue injury (AKI). The effect of LPS inclusive of EEG findings is recapitulated by peripheral or central injection of IL-1β or TNF, but not of IL-6^55,78^. Human studies also support the hypothesis that systemic inflammation causes central nervous effects and neuroinflammation. Endotoxin from salmonella cell walls injected into healthy volunteers dose-dependently affects cognitive function, emotional state and sleep^79,80^. LPS injection into healthy volunteers increased cytokine serum levels and sickness behaviour^81^, and an MRI study showed that LPS injection causes robust microglial activation in humans^81^.

Several pre-clinical studies also show evidence of acute neuroinflammation after AKI (bilateral renal ischemia-reperfusion-injury (IRI)), but neither how neuroinflammation was induced nor whether delirium was present was studied. In mice, at 2 and 24 hours after AKI the proinflammatory neutrophil chemoattractant C-X-C motif chemokine ligand 1 (CXCL1) and granulocyte colony-stimulating factor (G-CSF), but not IL-1β, TNF, IL-6, or CCL2 were upregulated in the cerebral cortex and hippocampus. Glial fibrillary acidic protein (GFAP) expression, a marker of astrocyte activation, increased in the cortex and corpus callosum. This was associated with decreased locomotor performance in the open field test^6^. Another study showed that AKI acutely increased serum CXCL1, G-CSF, and CCL2 levels, increased CXCL1, G-CSF, and CCL2 in the brain (but not IL-1β, TNF, IL-6), increased microglial numbers in the hippocampus and cortex, and induced transcriptional changes in the hippocampus^21^. A third AKI study reported upregulation of the damage-associated molecular pattern receptor Toll-like-receptor 4 (TLR4) in the hippocampus and striatum^22^. Except for monocyte-chemoattractant CCL2, we did not find mRNA upregulation of cytokines in the hippocampus or cortex (not shown). This may reflect our use of bulk mRNA, which could obscure cytokine expression in distinct low-abundance cell populations. Age is a well-known risk modifier for microglial responses to inflammatory challenge and for delirium^74,82^. Aged mice would thus likely show more prominent neuroinflammation, microglial activation and delirium-like symptoms. No previous AKI study reported hippocampal monocyte uptake. Yet rodent studies of sepsis-induced tissue injury have found that peripheral CCR2^+^ monocytes infiltrate the brain parenchyma, including the hippocampus, during early sepsis, with long-term microglial activation and proliferation^83,84^, both of which have been linked to sepsis-associated encephalopathy and delirium-like symptoms^56,57,85,86^.

Collectively, these studies largely support findings in our AKI model, with some exceptions. First, we did not find BBB leakiness after AKI at 6 hours and 48 hours timepoints. One study found BBB leakiness 24 hours after AKI using Evans Blue in a significantly more injurious AKI model (60 min of ischemia versus our 20-25 min)^6^. We did, however, find mild OPN-induced lung endothelial leakage in our AKI model^23,24^, suggesting that such mild leakage may also occur at the BBB; this could be ascertained by direct examination of brain vessels during AKI. Other systemic inflammatory stimuli, in particular sepsis, have also been linked to BBB disruption^87–89^. In humans, correlative but no direct evidence supports that elevated blood levels of the β-subunit of S100 protein (S100-β) reflect increased BBB permeability^90^, and serum S100-β increases in acutely ill elderly medical patients correlated with delirium^91^. However, no direct evidence for BBB disruption in human AKI or delirium exists, and some data question this hypothesis. For example, elevated cerebrospinal fluid (CSF) albumin reflects increased blood–CSF barrier permeability, yet CSF albumin was not increased in most patients with delirium, suggesting an intact BBB^92^. Second, we did not measure circulating classical cytokines and so cannot conclude whether they are involved or act up- or downstream of OPN. It is however well-documented in humans and in mice that AKI induces systemic inflammation with upregulation of serum IL-1β, TNF, IL-6, and CCL2, among other classical cytokines and chemokines (reviewed in^93^), and that this increases delirium risk in adults undergoing major surgery or who are critically ill^25,94–96^. As example, elevated plasma IL-1β correlates with delirium in patients with sepsis^97–99^. Like OPN^23,100–108^, these cytokines and chemokines are elevated in the serum of many inflammatory diseases or conditions associated with delirium, including trauma and surgery, various other sterile tissue injuries, and sepsis^25,56^, allowing the possibility that OPN has a broader role in delirium than after AKI. Risk factors, such as age and cytokine/chemokines serum level, may be key determinants of whether they trigger or contribute to AKI induced delirium.

Although osteopontin expressed in the brain or spinal cord is protective in direct brain or spinal cord injury^109–112^, our presented and published data^23,24^ show that circulating OPN released by the AKI kidney, in contrast, has detrimental effects on remote organs, in particular the lung^23^ and, as presented in this manuscript, the brain. Unpublished data in our AKI-induced remote lung inflammation model, and data presented here, show that the remote function of OPN is dependent on thrombin cleavage in the lung. The relevance of OPN cleavage by thrombin in vivo was first shown in cancer tissue^31,113^. We showed that both OPN fragments, OPN-N and OPN-C, were required to induce CCL2 in CCR2^+^ monocyte-derived peritoneal macrophages and endothelial cells. Thrombin cleavage of OPN endows OPN-N with enhanced ability to bind to leukocyte adhesion integrins (α4β1 and α9β1) as compared to OPN full-length^114–116;^ and liberates OPN-C to freely interact with CD44 variants^52,117–119^, suggesting they may be OPN fragment receptors. OPN can indeed activate focal adhesion kinase (FAK) or src-kinase and regulate integrin-linked kinase 1 (ILK1), as well as Akt and ERK signaling^120,121^, known integrin^122,123^ or CD44^124,125^ downstream effector molecules, respectively.

We had previously observed significant lung capillary thrombosis in our AKI-induced and OPN-dependent remote lung inflammation phenotype^23^ and had therefore suspected the lung as the site of thrombin activation and OPN cleavage. The large lung capillary bed is reached first by inflammatory signals released from the kidney (first pass), causing strong endothelial stress and activation, likely the stimulus needed to activate thrombin in the lung vasculature. Activating thrombin in the brain vasculature would risk thrombosis and catastrophic consequences, so OPN cleavage in the lung may confer an evolutionary survival advantage despite the risk of hypoxemia.

## Materials and Methods

### Animals and ischemia reperfusion injury

For all studies, adult (8 to 12 weeks old) male mice were used in accordance with the animal care and use protocol approved by the Institutional Animal Care and Use Committee of Washington University School of Medicine (IACUC 25-0049). C57BL/6J (B6) (JAX stock no. 000664) and B6.129S6(Cg)-Spp1tm1Blh/J (OPN−/−) (JAX stock no. 004936) were purchased from the Jackson Laboratory. OPNKI mice were generously gifted from Dr. Lawrence Leung, Stanford University School of Medicine. CCL2RFP^flox/flox^ mice CCR2^CreERT2tdTomato+/-^ mice were bred in our mouse facility. To initiate the activity of Cre recombinase and mark the CCR2 positive monocytes, mice were gavage with Tamoxifen (SIAL-T5648-1G, Sigma Aldrich) in corn oil (C8267, Sigma) p.o. at a dose of 10mg/Kg once on the day before sham/AKI surgery. Bilateral renal ischemia injury was performed as described previously^23,24^. Briefly, the renal pedicles were cross clamped for 18 or 20 min at 37°C in both kidneys using the flank approach by clamping both the kidney peduncles. Sham operations were performed with exposure of both kidneys but without induction of ischemia.

### OPN/ CCL2 neutralizing antibody injection

To assess whether OPN is necessary for the development of remote brain injury after acute kidney injury, we implemented OPN nAb experiment on *wt* mice. Mice were injected with either control Goat immunoglobulin G (IgG) or anti-OPN (AF808, R&D Systems) neutralizing antibody at the dose of 30 μg/mouse intraperitoneal two hours before the AKI surgery. For the CCL2 neutralizing experiment, mice injected control Hamster IgG or anti-CCL2 antibody at the dose of 200 μg/mouse (BE0185-R005MG, BioXCell). Mice underwent behavioral testing (EPM) and sacrificed at 48 hours after the AKI surgery.

### Behavioral tests

Animals underwent contextual fear conditioning tests at day1 and day2 after AKI followed by open field test (OFT) and elevated plus maze test (EPM) on day 2 following the previous protocols^126^. All testing was performed during the first 4 h of the light cycle by an experimenter blinded to group assignment.

Fear memory was analyzed using the fear conditioning test. At 48h, only the two day paradigm was done with a conditioning day at 24h after AKI and Contextual recall at 48h after AKI. On conditioning day 1, mice were placed in context A for 10.5 min and received 5 tone-shock pairings (30 s tone with 0.5 mA shock during the last 2 s). Immobility time in 30 seconds epochs was measured by a blinded observer. On day 2 (contextual test), mice were placed in context A for 10.5 min and immobility time was measured to assess contextual fear memory.

Anxiety-related and risk-taking behavior of mice was evaluated using EPM. When mice are placed in an EPM apparatus, both exploratory and fear drives are evoked, with rodents displaying a natural aversion for the open arms of the maze. Sham vs AKI Mice were evaluated at 48 hours, 1 month and 3 months after the surgery. The EPM consisted of two opposing open arms (35×5 cm) and two closed arms (35×5×15 cm) that extended from a central platform (5×5 cm) elevated 60 cm above the floor. A small, raised lip (0.5 cm) around the edges of the open arms prevents animals from slipping off. Mice were placed individually on the central platform facing an open arm, away from the examiner, and were allowed to freely explore the maze for 5 min under even overhead fluorescent lighting, as described previously .31 The behavior of each mouse was monitored using a video camera (Sony Exwave HAD; Sony Electronics) and the movement of mice was automatically registered and analyzed with a computerized tracking system (Viewer3; BIOBSERVE GmbH). Time spent in the open and closed arms was recorded, and each mouse was only tested once in the maze. The floor of the EPM was washed after each testing with 70% ethanol to remove odors left by previous subjects.

### Blood urea nitrogen assay

Renal function was measured with BUN levels at 24h and 48h after AKI from serum collected from sham and AKI mice using the DiUR100 kit (Thermo Fisher Scientific) according to the manufacturer’s instructions.

### Tissue processing

Mice were anesthetized by isoflurane overdose followed by transcardial perfusion with ice-cold 0.1 M heparinized phosphate-buffered saline (PBS, pH 7.4). Perfused brains were kept in 4% PFA at 4°C overnight, followed by equilibration in 30% sucrose for at least 48 h before sectioning using a freezing microtome at 30um thickness. For biochemical techniques, brains regions were rapidly removed, dissected, and flash frozen. For flow cytometry assays, brain tissues were collected in dissociation buffer on ice.

### Cell suspensions

The hippocampus and cortex tissues was obtained from one hemisphere of the brain. Number of samples used in each experiment are indicated in the respective figure legends. The hippocampal and cortex tissues were dissected on ice and digested at 37°C for 15 min with collagenase D (400 units/mL, Roche) in Dulbecco’s PBS (Lonza, Basel, Switzerland), each containing 50 μg/mL of DNase I (Sigma-Aldrich). The tissue was then mechanically dissociated with a glass Pasteur pipette, and centrifuged at 300g for 15 min. A 25% Percoll (Sigma-Aldrich) column was used to remove cell debris and myelin, followed by centrifugation at 1000 rpm for 10 min. Cell suspension was filtered through a 70 µm nylon filter before proceeding with staining. For immune cell isolation from blood, blood samples were collected into heparinized tubed (#365965 BD) at the time of sacrifice. 60µl of blood was added to 750 µl of RBC lysis buffer (#420302, Biolegend) and incubated in a rotator for 15 min at room temperature.

### Flow cytometry analysis

Cells were incubated for 5 min at RT with Zombie NIR Dye (1:2000, BioLegend, San Diego, CA, USA) to assess their viability. The Zombie NIR Dye was quenched, and cells were washed with cytometry buffer and blocked with FcR blocking reagent (1:50, Miltenyi Biotec, Bergisch Gladbach, Germany). Then, the samples were washed with cytometry buffer, stained with antibodies (Table 1) for 15 min at RT, and analyzed on a BD LSRFortessa flow cytometer (BD Biosciences, Franklin Lakes, NJ) using the Software v10.6.1 (BD Biosciences, Franklin Lakes, NJ). Cells were identified as (1) Microglia (CD45^int^ CD11b^+^), (2) neutrophils (CD45^hi^ CD11b^+^ Lineage^-^Ly6G^+^), (3) monocytes (CD45^hi^ CD11b^+^ Lineage^-^ Ly6G^-^ Ly6C^+^). Fluorescence minus one (FMO) and isotype control antibodies were used as negative controls for each marker.

**Table 1.** List of antibodies used in the experiments described in materials and methods.

| Antibody | Fluorophore | Species | Dilution | Source | Catalog number |
|---|---|---|---|---|---|
| CD45 | BV425 | Rat monoclonal | 1:500 | BioLegend | 103134 |
| CD11b | BV510 | Rat monoclonal | 1:500 | BioLegend | 101263 |
| Ly6C | BV785 | Rat monoclonal | 1:500 | BioLegend | 128041 |
| Ly6G | AF700 | Rat monoclonal | 1:100 | BioLegend | 108422 |
| CD68 | PE | Rat monoclonal | 1:100 | BioLegend | 137014 |
| Iba1 |  | Rabbit polyclonal | 1:500 | Fujifilm | 019-19741 |
| CD31 |  | Rat monoclonal | 1:200 | BD Pharmingen | 550274 |
| RFP |  | Goat polyclonal | 1:500 | Rockland | 200-101-379s |
| GFAP |  | Rabbit polyclonal | 1:500 | Dako | Z0334 |
| Neun |  | Rabbit monoclonal | 1:500 | Thermo fisher | 1362077 |
| TdTomato |  | Goat polyclonal | 1:500 | Sicgen | AB8181-200 |
| Secondary antibody | AF488 | Donkey anti-rabbit | 1:400 | Thermo Fisher | A-21206 |
|  | AF594 | Donkey anti-goat | 1:400 | Invitrogen | A-11058 |
|  | AF647 | Donkey anti-rat | 1:400 | Jackson ImmunoResearch | 712-545-153 |

### RNAscope-CCL2 in situ-hybridization (ISH)

To visualize the localization of CCL2 mRNA CCL2 in the brain ISH was performed following the manufacturer protocol (RNAscope™ Probe - Mm-CCL2 # 31179, Bio-Techne, Italy) on brain slices. Briefly, fixed brain sections were exposed to hydrogen peroxide and post-primary fixation using Neutral Buffered Formalin (Merck, Italy) for 12 min and then to the treatment with Protease Plus III (Biotechne, Italy). The treatment was followed by the ISH protocol using the CCL2 probe (#31179, Biotechne, Italy). After mounting on the microscopic slides, brain sections were imaged at hippocampal dentate gyrus region and prefrontal cortex regions using Nikon Eclipse E800 microscope.

### Histology, immunofluorescence staining, and quantification

In order to visualize CCL2 expressing cell types in the brain, we used CCL2RFP^flox/flox^ mice. For immunofluorescence, brain slices were permeabilized in 0.1% Triton X-100 for 3 min, and blocked for 1 hour with 10% normal goat serum supplemented with 1% bovine serum albumin (BSA) in PBS. Sections were incubated with primary antibodies in blocking solution overnight at 4°C. The primary antibodies used were goat anti-RFP (200-101-379s, Rockland; 1:500), rabbit anti-Iba1 (77973-870, Fujifilm; 1:500), rabbit anti-GFAP (Z0334, Dako; 1:500), rat anti-CD31 (550274, BD Pharmingen; 1:200). After extensive washes with PBS, fluorescently conjugated secondary antibodies were applied at 1:400 dilution for 1 hour at room temperature. The secondary antibodies used were as follows: Alexa Fluor 594 donkey anti-goat (A-11058), Alexa Fluor 488 donkey anti-rabbit (711-605-152), and Alexa Fluor 647 donkey anti-rat (712-545-153). Images were acquired with an Olympus FV1000 laser scanning confocal microscope at a ×200 magnification.

### Reverse transcription-quantitative polymerase chain reaction

RNA extraction, complementary DNA (cDNA) synthesis, and RT-qPCR were conducted in accordance with previous studies^127^. Total RNA was isolated from hippocampus and kidney tissue using Trizol (Invitrogen, Carlsbad, CA). cDNA was made using the QuantiTect® Reverse Transcription Kit (#205314, QIAGEN GmbH, QIAGEN Strasse 1, 40724 Hilden, Germany). The cDNA was diluted with RNase-free water to a final concentration of 8 ng/µL, and the samples were kept at −80 °C. RT-qPCR was carried out using Quantifast® SYBR® Green PCR Kit (#204054, QIAGEN GmbH, QIAGEN Strasse 1, 40724 Hilden, Germany) and Quanstudio TM 6Flex Real-Time PCR Systems (#4484642, Applied biosystems by Thermo Fisher Scientific) following the manufacturer’s instructions. Primers used are listed in the Table 2. The annealing temperature for the reaction was 58 °C, and the built-in software created the amplification curves and calculated the threshold cycle values. The GAPDH reference gene was used to normalize all the readouts. Data were reported as the mean relative values compared to the CON group using the 2−ΔΔCT method.

**Table 2.**
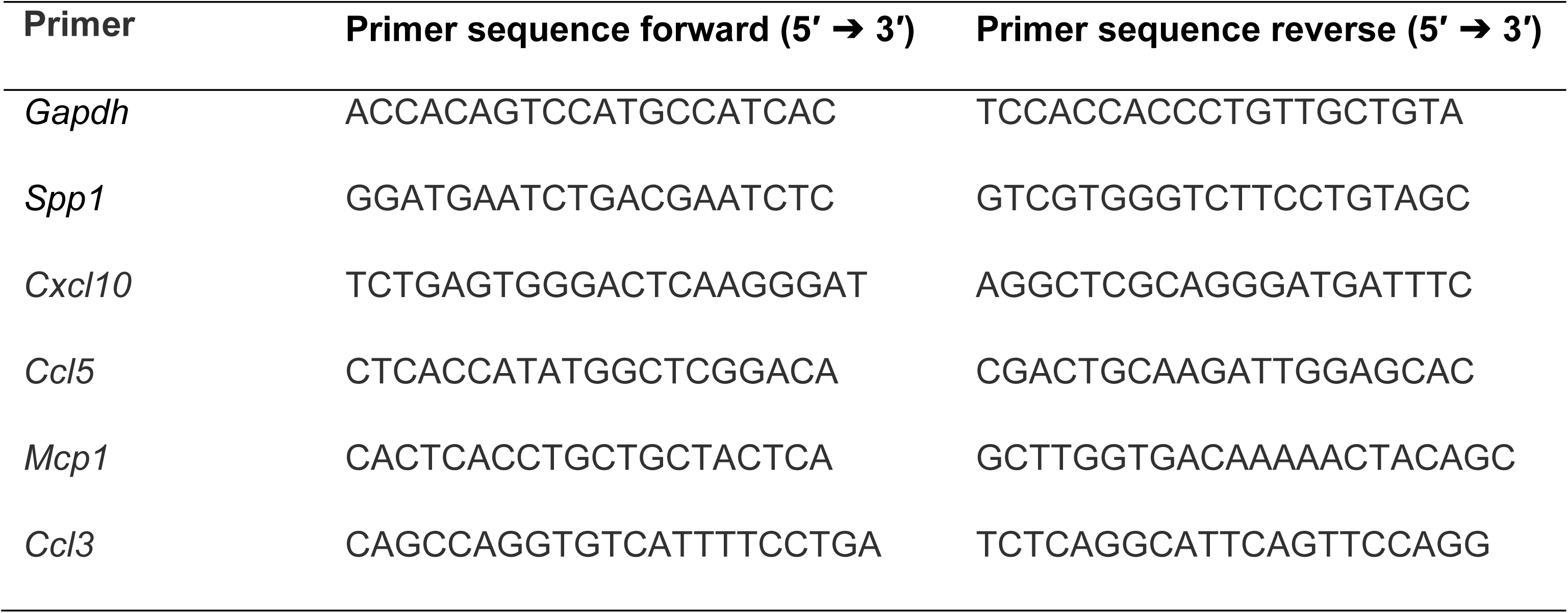
List of primers used in qPCR described in materials and methods.

### Three-dimensional reconstruction of microglia

We followed a published protocol for microglia morphology analysis^128^. 30-μm sections were stained with Iba1 (Table 1) 4°C overnight, followed by Alexa Fluor 488–conjugated secondary antibody (Table1) staining for 2 h. Sections were mounted on glass slides in TBS-X, dried, and coverslipped with mounting medium for fluorescence with DAPI. Imaging was performed on a Zeiss LSM 880 confocal laser scanning microscope (Zeiss, White Plains, NY) using a 20X 0.8 NA objective. Z-stacks were done with 1.00-μm steps in z direction; 1,024 × 1,024 pixel resolution were recorded and analyzed using IMARIS software (Bitplane, Concord, MA). Six microglial cells from the hippocampus *dentate gyrus* region were reconstructed and analyzed per mouse.

### Blood–brain–barrier assessment

BBB permeability was measured using fluorescein isothiocyanate (FITC)-dextran (Sigma-Aldrich). Two minutes prior to sacrifice, mice were given 50 µl of 70-kDA FITC-dextran (100 mg/mL) i.v. Brains were quickly removed after decapitation and fixed by immersion in 4% PFA for 24 h, followed by 48 h in 30% sucrose. Brains were embedded in cryostat compound (Tissue-Tek, Torrance, CA), frozen on dry ice, and cut into 50-μm thick sections on a cryostat. Z-stack images at 4 mm steps of the ipsilateral hippocampus were obtained at 20X with a Zeiss AxioImager Z2 (Zeiss) equipped with an ApoTome 2 optical sectioning grid imager. Image analysis was performed with ImageJ as previously described^128^.

### Enzyme-linked immunosorbent assay

Mouse OPN levels were measured in mouse serum, and tissue using enzyme-linked immunosorbent assay (ELISA) kits (DY1433 from R&D Systems) as per the manufacturer’s instructions. Serum dilutions for ELISAs was 1:4000 dilution, kidney lysates 1:4000 dilutions, and hippocampus 1:300 dilutions.

### Mass spectrometry for uremic retention solutes

Mass spectrometry on serum samples from sham/AKI mice was performed as previously described^129^. For data collection, metabolite extracts were run on either a Thermo Q-Exactive Orbitrap or an ID-X II Orbitrap Mass Spectrometer interfaced with a Thermo Vanquish Horizon LC system using an iHILIC-(P) Classic 2.1 mm × 100 mm, 5 μm column (HILICON, 160.102.0520) with an iHILIC-(P) Classic Guard column (HILICON, 160.122.0520) attached. The column temperature was set to 45 °C. The mobile phases were as follows: for solution A, 95% water, 5% acetonitrile, 20 mM ammonium bicarbonate, 0.1% ammonium hydroxide solution (25% in water), 5 μM medronic acid; for solution B: 95% acetonitrile, 5% water. Each sample was injected (6 μL of intracellular and 2 μL of medium metabolite extract) and eluted with a linear gradient: at 250 μL/min 0–1 min 90% B, 1–12 min 35% B, 12–12.5 min 25% B, 12.5–14.5 min 25% B, 14.5–15 min 90% B, 15–20min at 400 μL/min and 90% B, and 20–22 min at 250 μL/min and 90% B for re-equilibration. MS/MS fragmentation data were collected using targeted methods at 20 eV. Metabolites were identified based on matching accurate mass, retention time, and MS/MS data to authentic reference standards in our in-house library or from curated databases for level one to two identification confidence. Chromatograms for selected metabolites were manually integrated in Skyline Daily (version 22.2.1.425).

### Electroencephalogram (EEG) electrode placement

Placement of EEG electrodes and EEG monitoring were performed as previously described^130^. Anesthesia will be induced in *wt* mice in a clear plexi-glass chamber with isoflurane 0.5−2% using isoflurane air−gas vaporizer and then maintained through a mouse anesthesia mask fixed to a sterotaxic frame. Four over the head will be shaved and animals will be placed in the head holder of the stereotaxic frame. Four pilot holes (two bilateral holes 1 mm anterior to bregma and two bilateral holes 5 mm posterior to bregma, each 1.5 mm lateral to the sagittal suture) were tapped through the skull to the dura mater using a 23-gauge needle. A prefabricated EEG head mount (Pinnacle Technology, Inc. #8201-EEG, 2-channel EEG with common reference, bipolar montage) was attached on top of the skull with superglue and four stainless steel screws (Pinnacle Technology, cat. no. 8209) were threaded into the pilot holes. An additional wire electrode will be placed under the neck muscle to record EMG activity. Teflon−insulated wire will connect the electrodes to a small connector plug, which will be mounted on the skull with dental acrylic completely covering the exposed opening to the surrounding skin. Skin is also closed with tissue glue.

### EEG monitoring after AKI surgery

AKI surgeries were performed one week after the electrode placement to ensure the full recovery of mice. Following the AKI surgeries and full recovery of mice, on the same day, animals were transferred to a customized rodent video−EEG monitoring unit, where they will be monitored by digital video and EEG. The connector plug was attached to a flexible wire leading to an EEG amplifier head stage, allowing uninhibited movement. Animals had free access to water and food. Mice were monitored for 48-72 hours after AKI injury. This time period also overlap with the time of which our behavioral assessments were made.

### Peritoneal macrophages treatment

Mice were injected with 1 ml of aged thioglycolate i.p. per mouse and waited for 2 days prior to collecting peritoneal cells. At the day of collection, mice were injected with 1ml PBS i.p per mouse and anesthetized and sacrificed mouse with isoflurane. Massage abdomen for approximately 10-15 sec. Mouse was with abdomen up on paper towel swabbed the abdomen with 70% alcohol or isopropanol. A small incision in the center of the skin overlying the peritoneal wall was made and the peritoneal fluid was collected. Peritoneal fluid was centrifuged (300 x g; 3 min) and collected cell pellet. Usually about 2-4 million resident peritoneal cells can be recovered from one C57BL/6 mouse using this method and about 50% are peritoneal macrophages. Cell pellet from one mouse was resuspended in 1 ml of complete RPMI medium, cells were counted. Peritoneal cells were cultured in a 12-well plate, 2 million/well in 1 ml cRPMI at 37 °C with 5% CO_2_ for 6-18 h. During this time, peritoneal macrophages adhere to the plastic surface. The floating non-macrophages were then washed away by adding and aspirating 0.5 ml cRPMI medium twice. For OPN fragment treatment, cells were treated with FL, N, C, N+C, KI and for the inhibitor experiment N+C were either treated with CD44V6 and integrinα9 inhibitors. Cells and supernatant were harvested 24 hours after the treatment and analyzed for monocyte chemoattractant expression.

### Statistical analysis

All data were analyzed using GraphPad Prism 10.3.1 (La Jolla, CA). Data results are presented as mean ± standard error of the mean. Data were analyzed with student t-test or one-way analysis of variance with repeated measures, where necessary, followed by Tukey’s tests for multiple comparisons with a significance level of p < 0.05.

---

**Supplemental Fig. 1.**
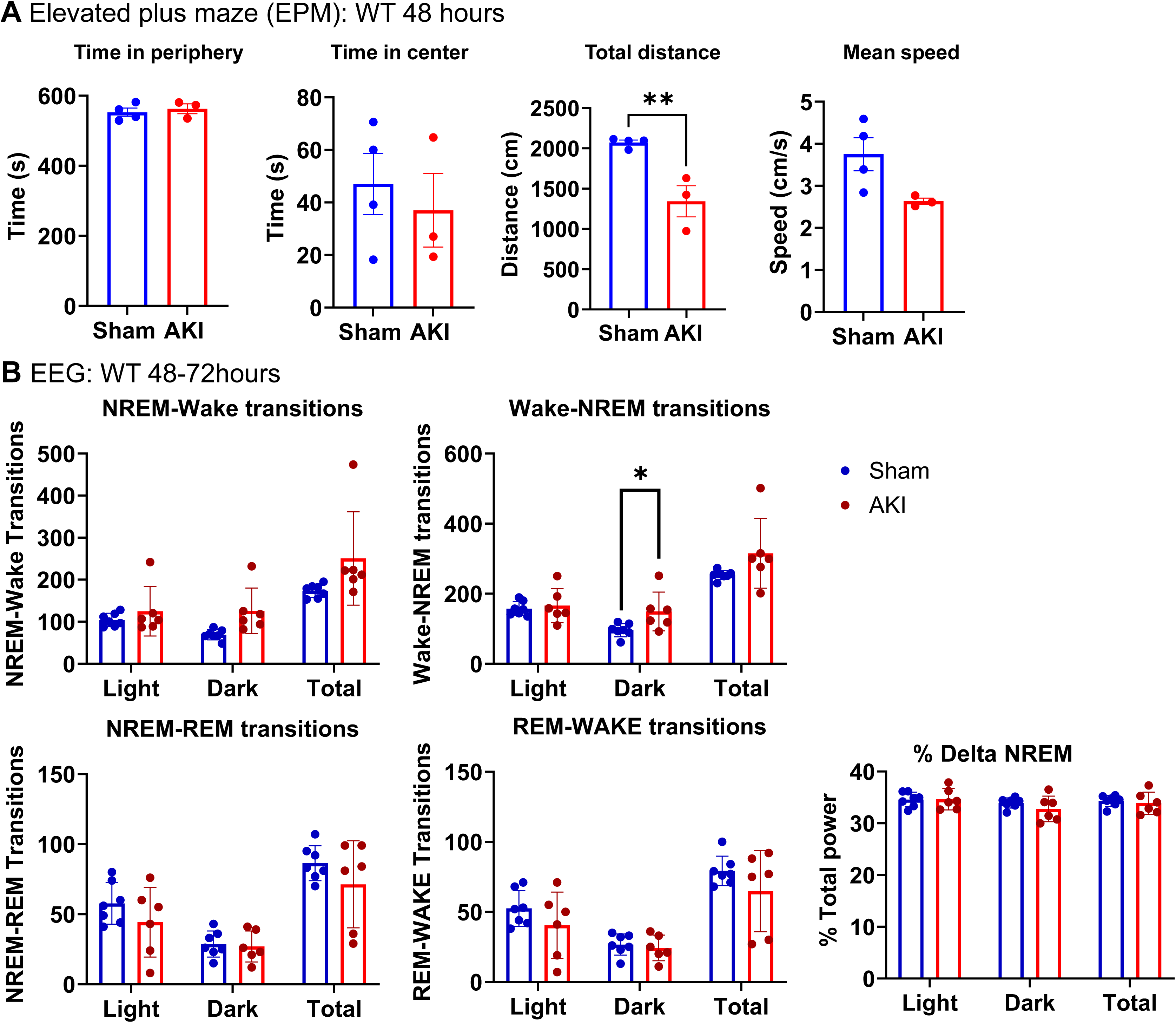
AKI induces acute neurological dysfunction with delirium-like symptoms in mice. (**A**) Additional parameters measured from the elevated plus maze test at 48h after sham/AKI in *wt* mice. (**B**) Additional parameters derived from from EEG recording done on *wt* mice at 48– 72 hours after sham/AKI. *n* = 7 to 10 animals per measurement. \**P* < 0.05 by student t-test. All data represent the mean ± SEM.

**Supplemental Fig. 2.**
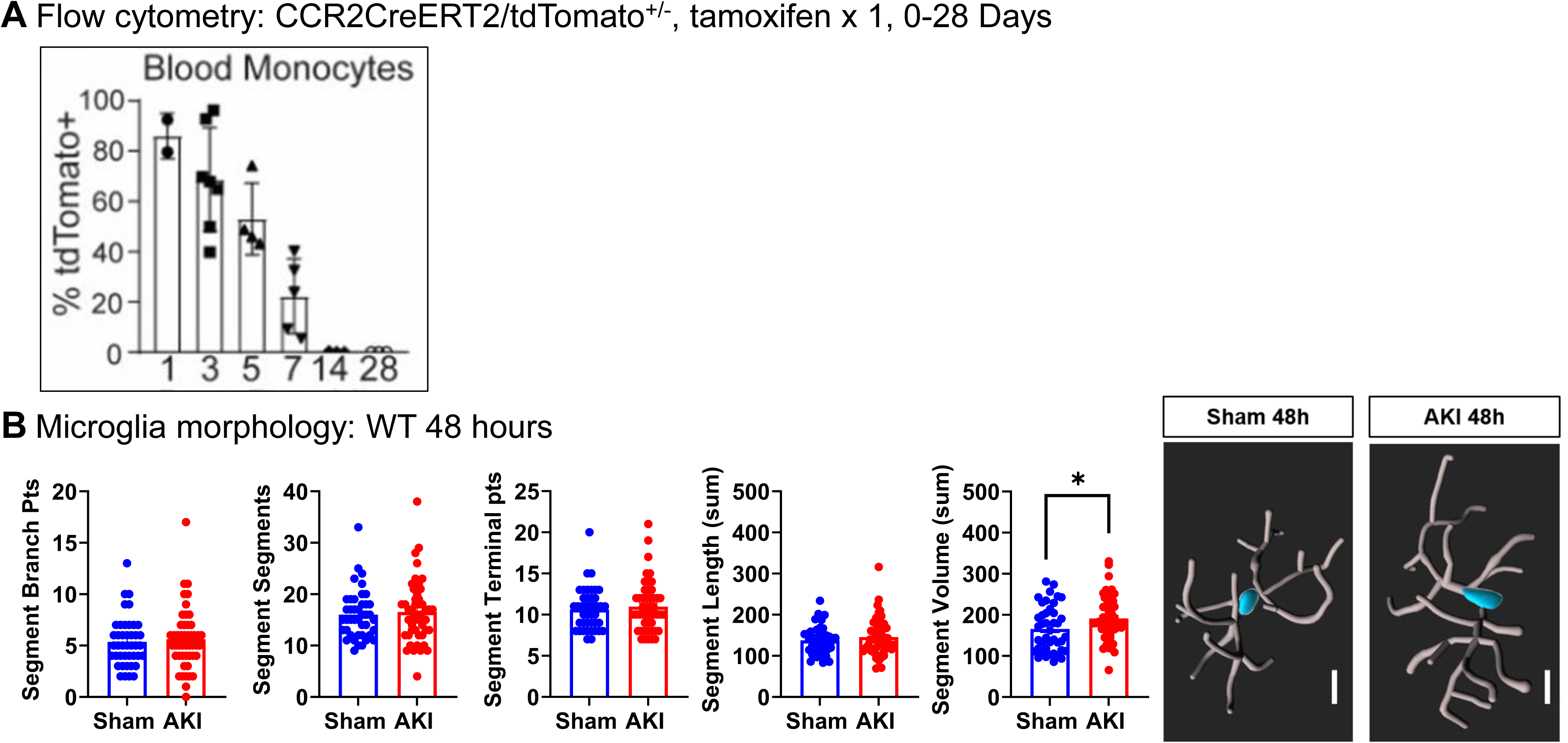
AKI induces acute hippocampal monocytic neuroinflammation without significant changes in microglial activation. (**A**) Flow cytometry of blood from CCR2^CreERT2tdtomato+/-^ mice 0-5 days. (**B**) Microglia three-dimensional reconstruction and morphology analysis (*n* = 6 to 10 animals per measurement. \**P* < 0.05, \*\**P* < 0.01, \*\*\**P* < 0.001 by student t-test. All data represent the mean ± SEM.

**Supplemental Fig. 3.**
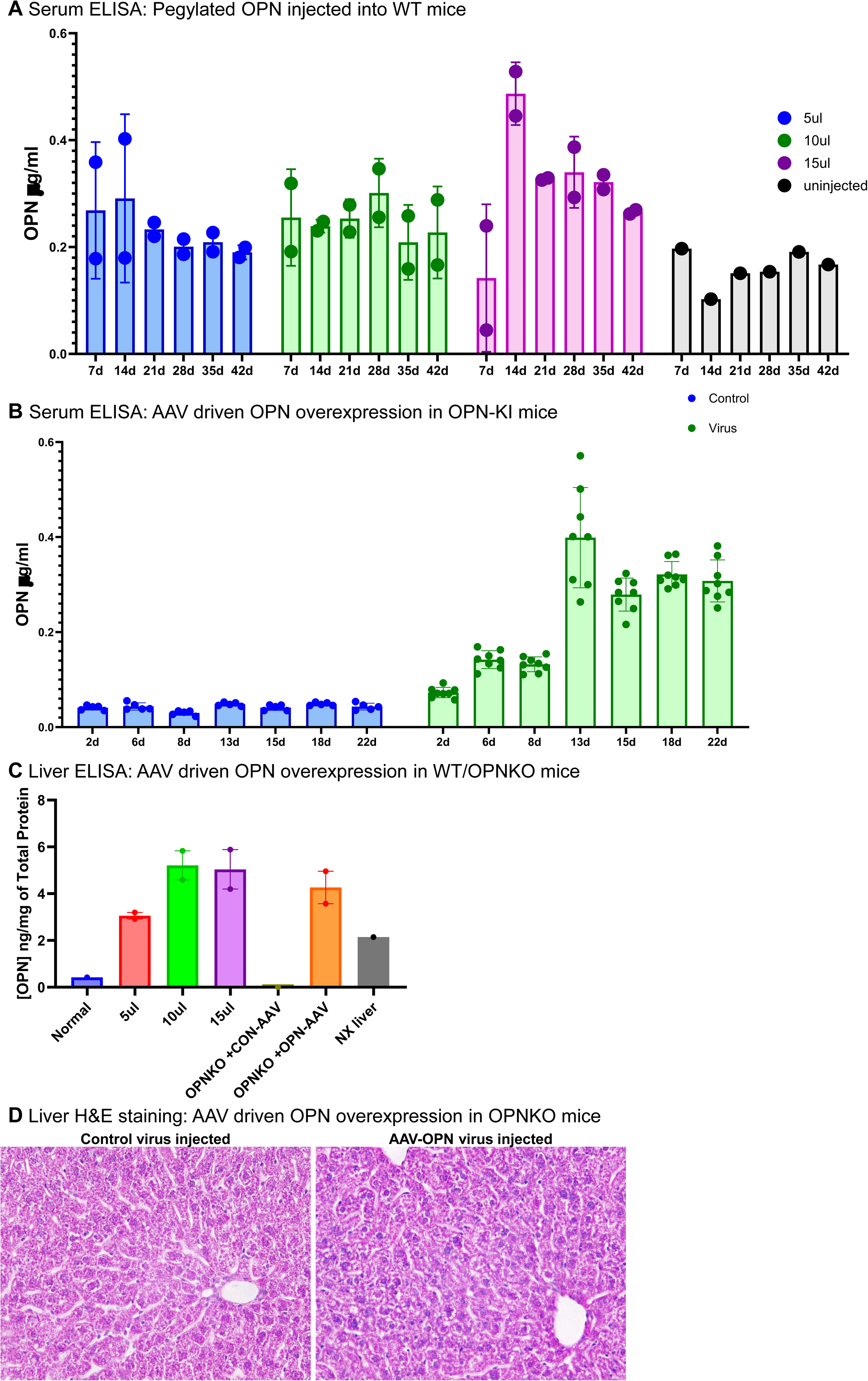
Attempts to overexpress OPN in *wt*, OPN-KO and OPN-KI mice. (**A**) Serum OPN levels over 42 days measured with ELISA in *wt* mice injected with pegylated OPN at different doses; 5, 10, and 15 µl/mouse. (**B**) Serum OPN levels measured with ELISA in OPN-KI mice with AAV8-driven liver overexpression of OPN at different time points (*n* = 2 to 7 animals per measurement. (**C**) OPN protein levels in the livers of OPN-KI mice injected with AAV-driven OPN to overexpress OPN at 22 days after the injection. (**D**) H&E staining of livers from AAV-driven OPN overexpression in OPN-KO mice showing pyknosis, condensed chromatin in the nuclei of hepatocytes in the AAV-OPN livers. (*n* = 2 to 7 animals per measurement). All data represent the mean ± SEM.

**Supplemental Fig. 4.**
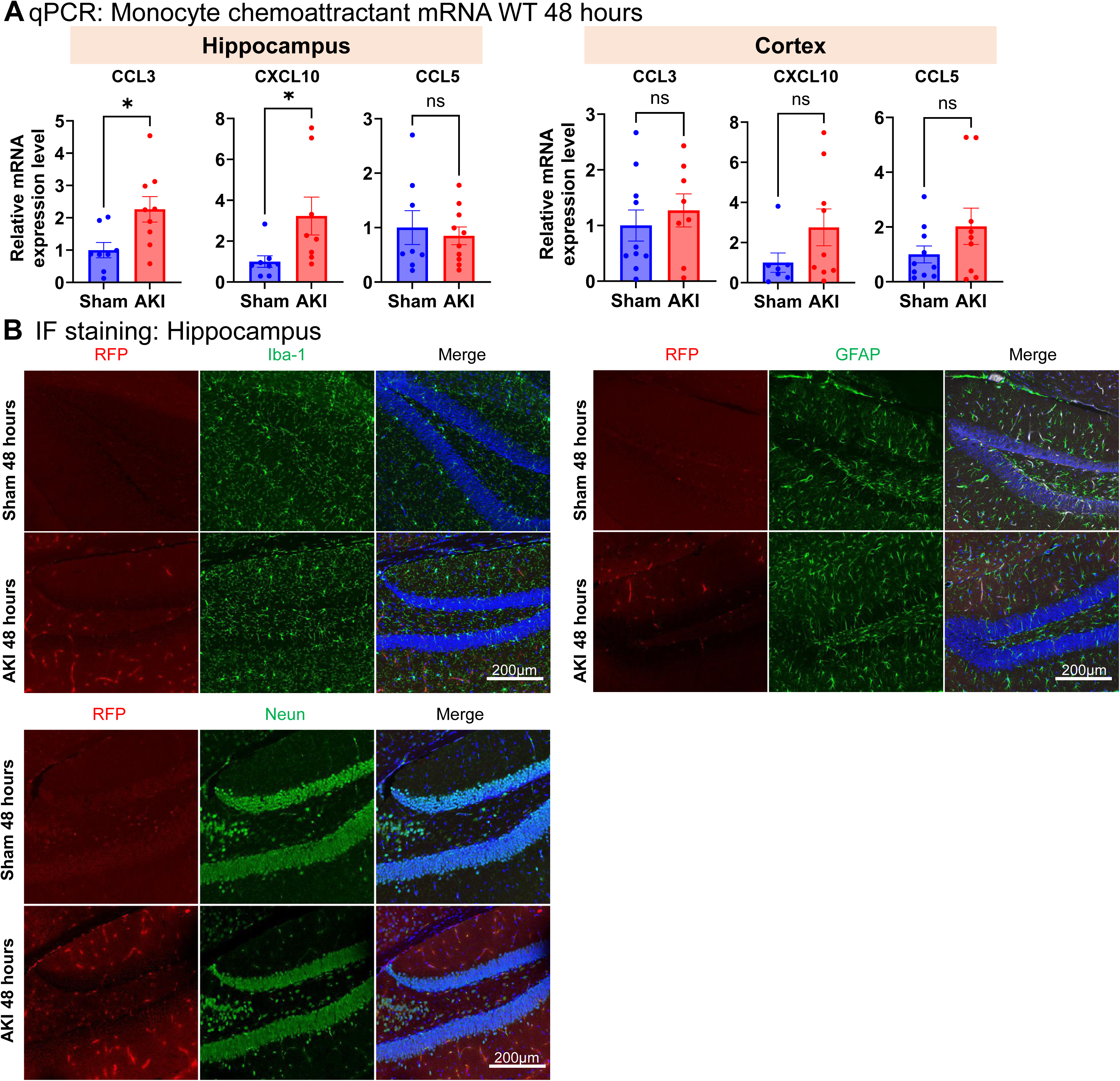
Expression levels of other monocyte chemo-attractants and CCL2 co-staining with cellular markers. (**A**) Monocyte chemoattractants *ccl3, cxcl10, and ccl5* mRNA expression levels assessed by qPCR in hippocampus and cortex at 48h after sham/AKI in *wt* mice. (**B**) Immunofluorescence staining of brain tissue showing RFP signal does not colocalize with Iba1 (marker for microglial cells) or Neun (Neuronal marker) in the hippocampus of CCL2-RFP^flox/flox^ mice. (*n* = 6 to 8 animals per measurement. \**P* < 0.05 by student t-test. All data represent the mean ± SEM.

**Supplemental Fig. 5.**
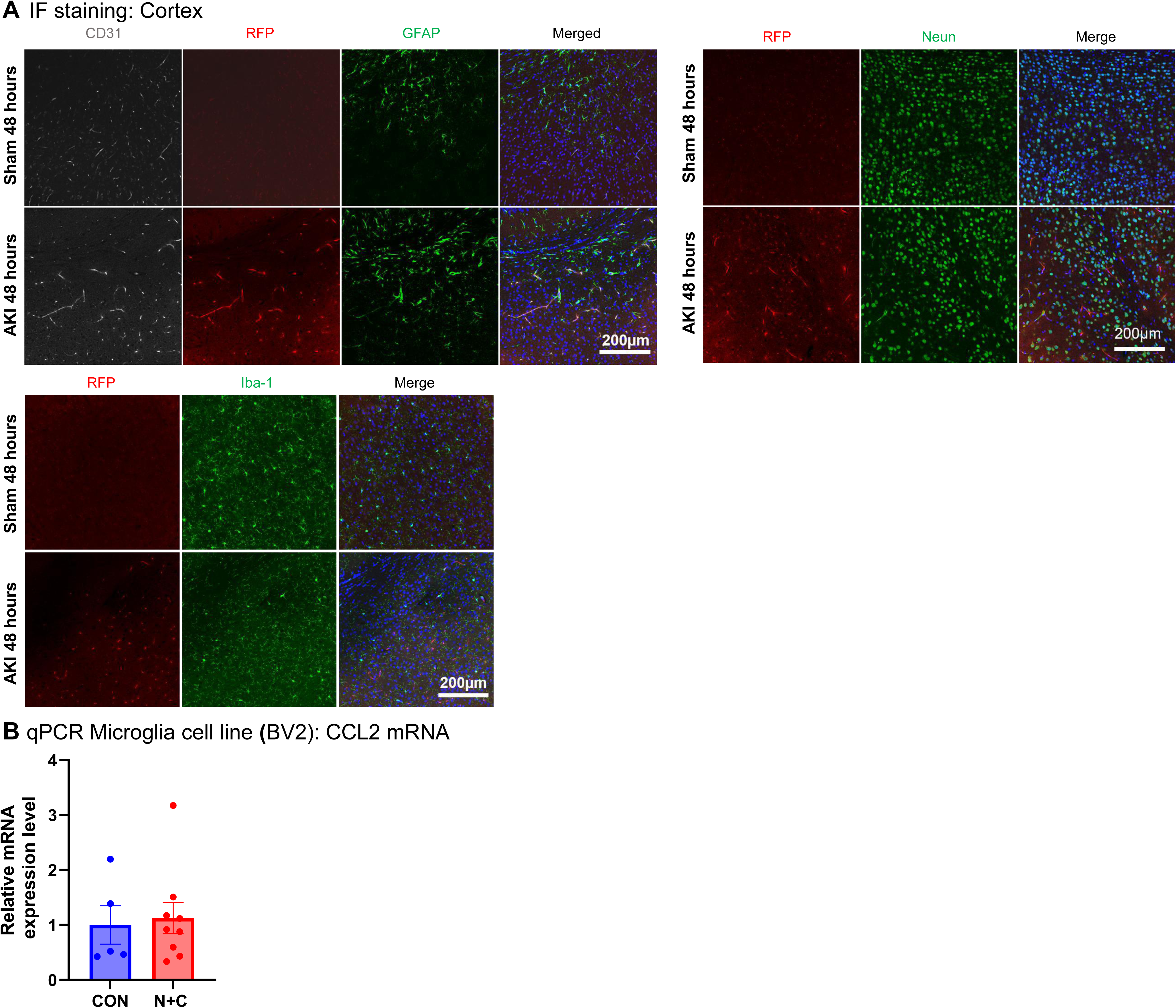
Colocalization of RFP signal with endothelial marker CD31 in the cortex, but not other brain resident cell types. (**A**) Immunofluorescence staining of brain tissue showing RFP signal colocalizes with CD31 (marker for endothelial cells) in the cortex regions of CCL2-RFP^flox/flox^ mice. RFP signal does not colocalize with GFAP (astrocyte marker), Iba1 (microglial marker), or Neun (neuronal marker). (**B**) Microglial cells (BV2) treated with either full length OPN or N+C OPN fragments do not show an increase of CCL2 mRNA (qPCR).

